# Heat Shock Factor 1 (HSF1) specifically potentiates c-MYC-mediated transcription independently of the canonical heat-shock response

**DOI:** 10.1101/2022.02.22.481519

**Authors:** Meng Xu, Ling Lin, Kun-Han Chuang, Babul Ram, Siyuan Dai, Kuo-Hui Su, Zijian Tang, Chengkai Dai

## Abstract

Despite its pivotal roles in biology, how the transcriptional activity of c-MYC is attuned quantitatively remain poorly defined. Here, we show that heat shock factor 1 (HSF1), the master transcriptional regulator of the heat-shock, or proteotoxic stress, response, acts as a key modifier of the c-MYC-mediated transcription. HSF1 deficiency diminishes c-MYC DNA binding and dampens its transcriptional activity genome-widely. Mechanistically, c-MYC, MAX, and HSF1 assemble into a transcription factor complex on genomic DNAs and, surprisingly, the DNA binding of HSF1 is dispensable. Instead, HSF1 physically recruits the histone acetyltransferase GCN5, thereby promoting histone acetylation and augmenting c-MYC transcriptional activity. Thus, our studies reveal that HSF1 specifically potentiates the c-MYC-mediated transcription, distinct from its role in the canonical heat-shock response. Importantly, this mechanism of action engenders two distinct c-MYC activation states, primary and advanced, which may be important to accommodate diverse physiological and pathological conditions.

## INTRODUCTION

The *MYC* proto-oncogene family encodes a class of bHLH/ZIP transcription factors consisting of C-, L-, and N-MYC, which govern a plethora of cellular functions including cell proliferation, differentiation, apoptosis, metabolism, and others^1, 2^. The most prominent member of this family is *c-MYC*. Dysregulation of *c-MYC*, occurring in over 70% of all human cancers, is associated with poor patient outcomes^3, 4^. Moreover, *c-MYC* is a key player in pluripotency reprogramming^5, 6^. Following heterodimerization with MYC-associated factor X (MAX), c-MYC binds to the E-box (5’-CACGTG-3’) element or its variants on genomic DNAs and regulates the transcription of up to 15% of all human genes^1–4^. To achieve effective DNA binding and transcription initiation, cofactors are recruited to remodel the chromatin architecture. Among these cofactors is the STAGA (SPT3-TAF(II)31-GCN5L acetylase) complex^7, 8^. Within this complex, GCN5/KAT2A is a histone acetyltransferase that can acetylate histone H3 at lysine 9 (H3K9), lysine 14 (H3K14), and other lysine residues^9, 10^. Histone acetylation facilitates the rearrangement of chromatins from a condensed state to a transcriptionally accessible state, permitting transcription factors to access DNA for gene expression regulation^11^.

Heat shock factor 1 (HSF1) is the master regulator of the heat-shock, or proteotoxic stress, response (HSR/PSR), an evolutionarily conserved cytoprotective transcriptional program helping cells adapt to a wide variety of environmental and pathological challenges^12, 13^. Following trimerization, nuclear translocation, posttranslational modifications, and recognition of the heat shock element (HSE), which is canonically composed of 5’-GAANNTTC-3’ nucleotide sequence motif^12, 13^, HSF1 governs the transcription of genes involved in protein folding and degradation, particularly molecular chaperones or heat shock proteins (*HSPs*), in response to proteotoxic stress. Contrasting with its broadly acclaimed role in maintaining proteomic stability and promoting survival under stress, HSF1 potently enables malignancy^14, 15^. The pro-oncogenic mechanisms of HSF1 appear to be multifaceted, including suppressing proteomic instability, impeding senescence and apoptosis, reprogramming metabolism, and even promoting immune evasion^16–21^. Whereas deletion of *c-Myc* in mouse embryos caused severe developmental defects in a broad range of organs^22^, *Hsf1* appears dispensable for embryonic development and cell viability in the absence of proteotoxic stress^23^. However, in stark contrast to their non-transformed counterparts, cancerous cells rely on HSF1 for their growth and survival, rendering it essential to malignancy^24^. Despite their importance to oncogenesis, whether there is an interplay between c-MYC and HSF1 remains unclear.

We herein report that HSF1 specifically potentiates the c-MYC-mediated transcriptional program. Mechanistically, HSF1, c-MYC/MAX dimers, and GCN5 constitute a previously unrecognized transcription factor complex, the assembly of which is fostered by c-MYC DNA binding. Through physical interactions with both partners, HSF1 recruits GCN5 to c-MYC, heightening histone H3 acetylation at c-MYC target gene loci, promoting c-MYC/MAX DNA binding, and, ultimately, augmenting transcriptional activity. Thus, our studies reveal a new mode of regulation through which HSF1 dictates the transcriptional capacity of c-MYC.

## RESULTS

### HSF1 is required for robust c-MYC transcriptional activity

Both c-*MYC* and *HSF1* are located on human chromosome 8q24.21-24.3, an amplicon frequently found in human cancers^25, 26^. According to the Cancer Genome Atlas (TCGA) PanCancer studies, amplification of *c-MYC* and *HSF1* occurs at 8% and 6% of patients, respectively. Among those patients with *c-MYC* amplification, approximately 59% display co-duplication of *HSF1* (co-occurrence, p<0.001, Fisher’s exact test) (Figure 1A). Moreover, the mRNA levels of *c-MYC* and *HSF1* are positively correlated in human cancers (Figure 1B). Given their prominent roles in malignancy, we reasoned that the co-amplification and co-expression of *c-MYC* and *HSF1* might be selected for oncogenesis. First, we set out to explore whether HSF1 impacts c-MYC transcriptional activity using a dual reporter assay, where the expression of secreted alkaline phosphatase (SEAP) is controlled by binding of c-MYC/MAX to the E-box elements fused to the minimal TATA-like promoter.

**Figure 1.**
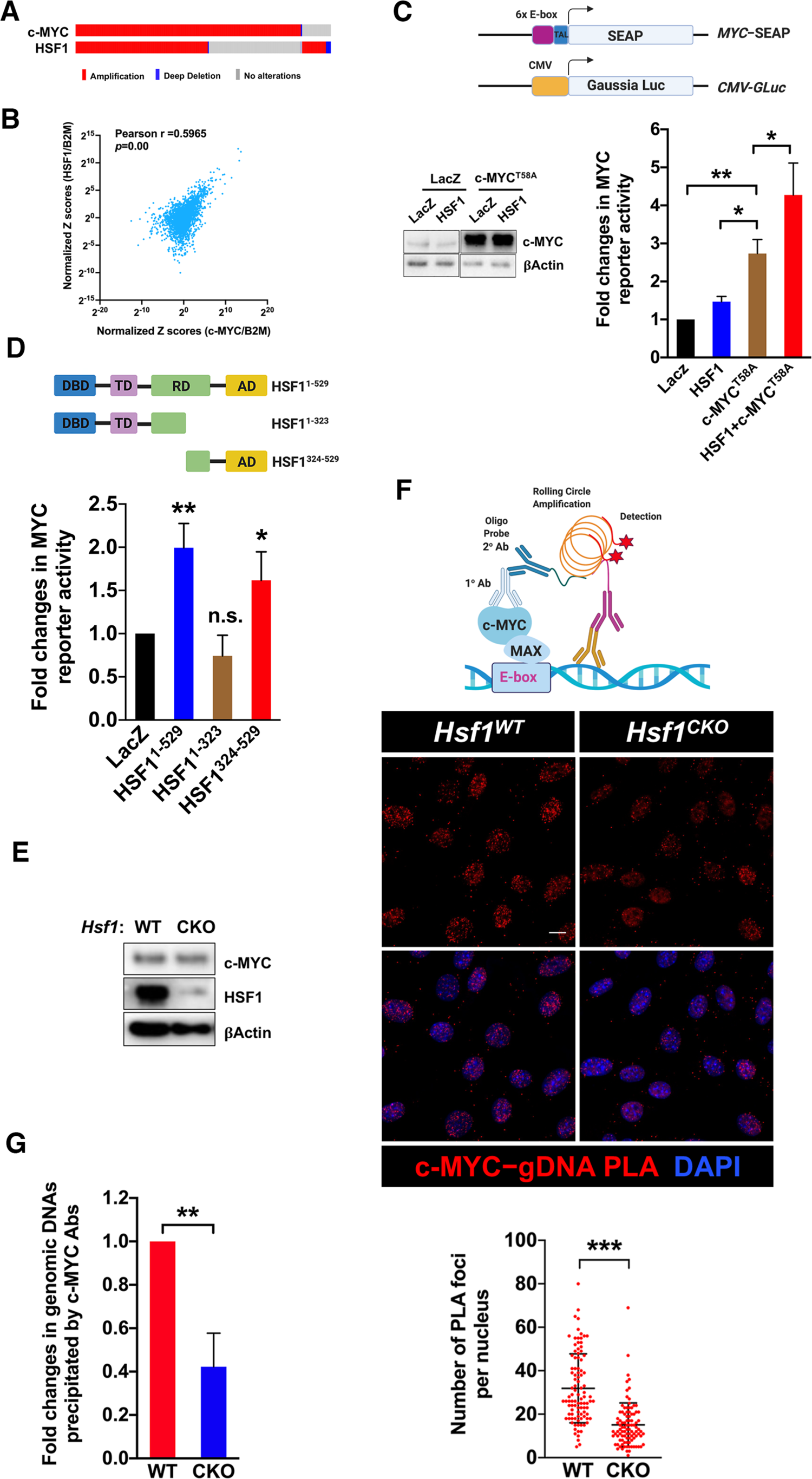
HSF1 is required for robust c-MYC transcriptional activity. (A) Co-amplification of *c-MYC* and *HSF1* in human cancers. Data are generated by the TGCA Research Network (https://www.cancer.gov/tcga). (B) Positive correlation between *c-MYC* and *HSF1* mRNA levels in human cancers. Analyses were performed using the GEPIA2 web server^27^. B2M: β-2-microglobulin. (C) The dual MYC reporter system, comprising an E-box element-driven SEAP plasmid and a CMV-driven Gaussia luciferase (GLuc) plasmid, were co-transfected with indicated plasmids into HEK293T cells for 48 hr (mean ± SD, n =3 independent experiments, One-way ANOVA). Cell lysates were immunoblotted. (D) Endogenous c-MYC activities were measured by the dual reporter system in HEK293T cells co-transfected with indicated plasmids (mean ± SD, n =3 independent experiments, One-way ANOVA). (E) *Hsf1* was deleted in immortalized *Rosa26-CreERT^2^; Hsf1^fl/fl^* MEFs treated with and without 4-OHT for 7 days. c-MYC levels were detected by immunoblotting. (F) Top panel: schematic depiction of c-MYC-gDNA PLA technique. Middle panel: visualization of endogenous c-MYC binding to genomic DNAs by PLA (red) in immortalized *Rosa26-CreERT2; Hsf1^fl/fl^* MEFs. Scale bars: 10µm. Lower panel: quantitation of c-MYC-gDNA binding by counting the numbers of PLA foci per nucleus (mean ± SD, n=98 nuclei, Mann Whitney test). (G) Quantitation of c-MYC-bound genomic DNA fragments following ChIP in immortalized MEFs (mean ± SD, n = 3 independent experiments, two-tailed Student’s *t* test).

Transient overexpression of *c-MYC^T58A^*, a mutant resistant to proteasomal degradation^28^, activated the reporter, as expected; of note, co-expression of HSF1 enhanced this activation (Figure 1C). HSF1 did not elevate the levels of endogenous or exogenous c-MYC proteins (Figure 1C), pinpointing a specific effect on c-MYC transcriptional activation. To demonstrate this c-MYC activation by HSF1 under physiological conditions, we examined the expression of several well-defined c-MYC target genes in immortalized mouse embryonic fibroblasts (MEFs) following *Hsf1* knockdown (KD). Considering that HSF1 becomes constitutively active in malignant cells, rendering them addicted to HSF1^14^, we elected to perform this experiment using this non-transformed cell type, for which HSF1 is dispensable. Two independent *Hsf1*-targeting siRNAs both diminished the transcripts of these target genes (Figure S1A).

Next, we asked whether this c-MYC activation requires the HSF1-mediated transcription. To address this, we expressed two mutants, HSF1^1–3^^23^ lacking the C-terminal transactivation domain (AD) and HSF1^324–529^ lacking the N-terminal DNA-binding domain (DBD), in HEK293T cells. Both mutants are deficient for transcriptional activity, as shown previously^20^. Interestingly, HSF1^324–529^, but not HSF1^1–323^, was sufficient to activate the c-MYC reporter (Figure 1D), strongly suggesting a transcription-independent mechanism. HSP90AA1/HSP90*α*, a transcriptional target of HSF1, was previously reported to stabilize c-MYC proteins^29^. Thus, it remains possible that HSF1 could regulate c-MYC via HSP90. However, HSP90 overexpression failed to rescue the diminished mRNAs of c-MYC target genes in *Hsf1*-deficient MEFs, despite elevated c-MYC proteins (Figure S1B and S1C), arguing against a direct activation of c-MYC by HSP90. Together, these results illustrate the necessity of HSF1 for c-MYC-mediated transcription and further indicate that HSF1 regulates c-MYC independently of its intrinsic transcriptional action.

### HSF1 promotes c-MYC binding to genomic DNAs

How does HSF1 affect c-MYC transcriptional activity? Unexpectedly, HSF1 impacted the DNA binding capability of c-MYC. This was detected by proximity ligation assay (PLA), a technique previously adapted to visualize interactions between transcription factors and genomic DNAs (gDNAs) *in situ*^30^. While the specificity of anti-dsDNA antibodies was demonstrated previously^30^, siRNA-mediated KD validated the specificity of anti-c-MYC antibodies (Figure S1D). Compared with *Hsf1* wildtype (WT) cells, PLA foci denoting the c-MYC-gDNA interaction were diminished in *Hsf1* conditional knockout (CKO) MEFs (Figures 1E and 1F), in which *Hsf1* deletion was induced by 4-hydroxytamoxifen (4-OHT)^31^. Importantly, this defect in DNA binding was confirmed by conventional c-MYC ChIP. When using equal amounts of chromatins, c-MYC antibodies precipitated less genomic DNA fragments from *Hsf1^CKO^* MEFs (Figure 1G).

To elucidate how broad this impact on DNA binding was, we employed the CUT&RUN-seq technique^32^, a new alternative to ChIP-seq, to profile genome-wide c-MYC DNA binding in these MEFs. Similarly, when using equal numbers of cells, less amounts of nuclease-digested DNA fragments were released from *Hsf1^CKO^* MEFs (Figure 2A). To account for this global change in c-MYC DNA binding, we spiked these released DNA fragments with equal amounts of *E. coli* DNAs as the normalization control. Following spike-in normalization, CUT&RUN-seq analyses revealed a genome-wide reduction in c-MYC DNA binding in *Hsf1^CKO^* MEFs (Figure 2B). Owing to the extremely low background signals, CUT&RUN-seq identified more than 200,000 binding sites in *Hsf1^WT^* cells; nonetheless, nearly 91% of these binding sites were located at either intergenic, intronic, or exonic regions (Figure 2C and Table S1). It has been known that c-MYC frequently binds to intergenic regions^33^. By contrast, approximately 70% of all binding sites identified in *Hsf1^CKO^* MEFs were associated with promoters, despite considerably diminished total binding sites (Figure 2C and Table S2). This finding indicates that *Hsf1* deficiency mostly abolished the c-MYC binding to non-promoter regions. Apart from this differential genomic distribution, binding sites in *Hsf1^WT^* cells displayed higher signals, a measure of c-MYC binding affinity, than binding sites in *Hsf1^CKO^* cells, especially those associated with promoters (Figure 2D). Within the same cell types, binding sites located in promoters displayed the highest signals; by contrast, those located at intergenic and intronic regions showed the lowest (Figure S2A).

**Figure 2:**
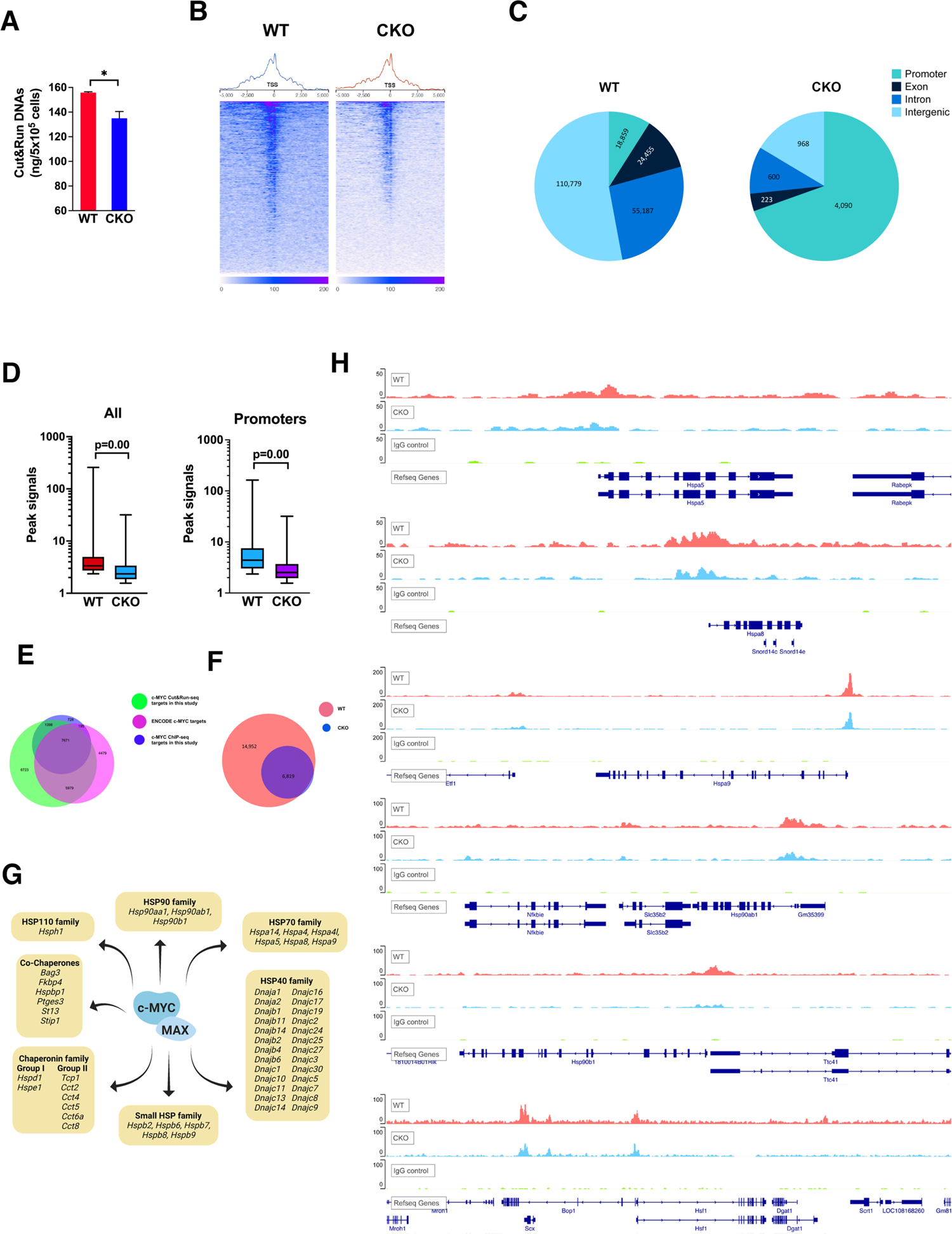
HSF1 promotes c-MYC DNA binding. (A) Quantitation of released genomic DNA fragments in the CUT&RUN experiments in immortalized MEFs (mean ± SD, n =2 biological replicates, two-tailed Student’s *t* test). (B) TSS plots of aligned CUT&RUN-seq reads following spike-in normalization (two biological replicates are combined). (C) Genomic distributions of CUT&RUN-seq peaks in *Hsf1^WT^* and *Hsf1^CKO^* MEFs. (D) Box plots of peak signals in *Hsf1^WT^* and *Hsf1^CKO^* MEFs. The box bounds the IQR divided by the median and the whiskers extend to the minimum and maximum values (Mann-Whitney U test). Left: all peaks (n=209,466 WT and 5,900 CKO); Right: peaks within promoters (n=18,859 WT and 4,090 CKO). (E) Venn diagram showing the overlaps of c-MYC target genes among different experiments. (F) Venn diagram showing the overlaps of c-MYC target genes identified by CUT&RUN-seq between *Hsf1^WT^* and *Hsf1^CKO^* MEFs. (G) Summary of c-MYC target genes encoding chaperones and co-chaperones. (H) Visualization of c-MYC binding to *Hsp* and *Hsf1* genes

Whereas commonly applied to histone modification studies, only a few transcription factors have been investigated using the CUT&RUN-seq technique. To demonstrate the validity of this new technique, we also performed the conventional ChIP-seq experiments using the very same antibody and *Hsf1^WT^* MEFs. While CUT&RUN-seq identified total 21,771 unique genes bound by c-MYC, ChIP-seq only identified 9,992 (Table S3). Of note, nearly 91% of those 9,992 genes were also detected by CUT&RUN-seq (Figure 2E), demonstrating a high degree of comparability between these two techniques. Importantly, our CUT&RUN-seq also identified 74% of ENCODE MYC target genes (18,324) (Figure 2E), considering the distinct experimental conditions. Moreover, CUT&RUN-seq peak sequences were highly enriched for the E-box motif; by contrast, the HSE motif was far less enriched (Figure S2B). In addition, peak visualization confirmed the binding of c-MYC to several classic target genes, including *Npm1*, *Ncl*, *Odc1*, *Cdk4*, and *Hspd1* (Figure S2C). Together, these results validate our CUT&RUN-seq experiments.

As expected, the c-MYC target genes in *Hsf1^WT^* and *Hsf1^CKO^* cells almost completely overlapped, although in *Hsf1^CKO^* cells c-MYC only bound to 31.8% of total target genes (Figure 2F). Despite weak signals in general, peak visualization confirmed the c-MYC binding to intergenic regions, (Figure S2D). Of note, an array of *Hsp* genes, spanning all HSP families, were identified as the targets of c-MYC (Figure 2G and 2H). Among them are several prominent constitutively expressed *Hsp* genes, including *Hspa5/Bip, Hspa8*/*Hsc70*, *Hspa9/Grp75*, *Hsp90ab1/Hsp84*, and *Hsp90b1/Grp94*. Of great interest, *Hsf1* was also a target of c-MYC (Figure 2H), a finding further confirmed by ChIP-seq (Figure S2E). These results support an important role of c-MYC in controlling cellular chaperoning capacity, both constitutive and inducible. Collectively, our findings indicate that HSF1 promotes c-MYC DNA binding genome-widely, a step crucial to its transcriptional activity.

### HSF1 physically interacts with c-MYC/MAX dimers

Prompted by the observation that the transcriptional activity of HSF1 is dispensable for c-MYC regulation, we next explored their potential physical interactions. Co-immunoprecipitation (co-IP) experiments in HEK293T cells revealed that exogenously expressed FLAG-HSF1 interacted with both HA-c-MYC and V5-MAX (Figure 3A). Importantly, this interaction also occurred under physiological conditions. PLA clearly detected the interaction between endogenous HSF1 and c-MYC, predominantly localized within the nucleus, in HeLa cells (Figure 3B).

**Figure 3.**
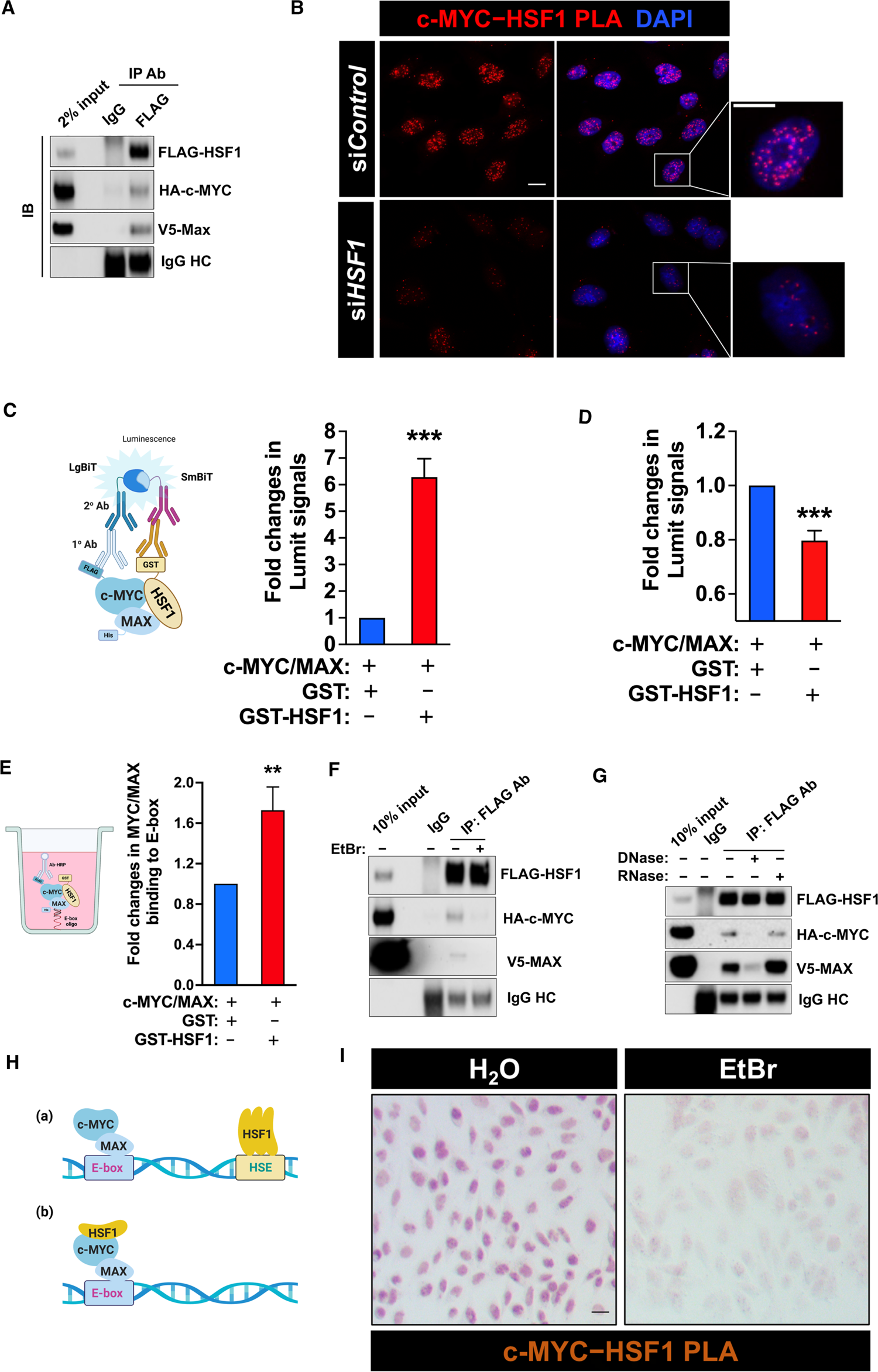
HSF1 physically interacts with c-MYC. (A) Co-IP of FLAG-HSF1, HA-c-MYC, and V5-MAX from transfected HEK293T cells (representative images of three independent experiments). HC: heavy chain. (B) Endogenous c-MYC-HSF1 interactions were detected by PLA in HeLa cells using a rabbit anti-c-MYC (D3N8F) antibody and a mouse monoclonal anti-HSF1 (E-4) antibody. Scale bars, 10µm. (C) *In vitro* direct interactions between recombinant HSF1 and c-MYC/MAX dimers were detected by the Lumit™ immunoassay. The reactions without primary antibodies were set up as the blanks, which were subtracted (mean ± SD, n =3 independent experiments, two-tailed Student’s t test). (D) *In vitro* interactions between recombinant c-MYC and MAX proteins, with and without recombinant HSF1 proteins, were detected by the Lumit™ immunoassay (mean ± SD, n =3 independent experiments, two-tailed Student’s t test). (E) *In vitro* binding of recombinant c-MYC/MAX dimers to E-box oligos, with and without recombinant HSF1 proteins, was detected by ELISA (mean ± SD, n =3 independent experiments, two-tailed Student’s t test). (F) Lysates of HEK293T cells co-transfected with indicated plasmids for 3 days were treated with EtBr (400 µg/mL) on ice for 30 min. The interaction of FLAG-HSF1 with HA-c-MYC/V5-MAX was detected by co-IP (representative images of three independent experiments). (G) Lysates of HEK293T cells co-transfected with indicated plasmids for 3 days were treated with either 10 U of DNase I or RNase at 37 °C for 20 min, followed by co-IP (representative images of three independent experiments). (H) Schematic depiction of two possible models of DNA-dependent protein-protein interactions. (I) Endogenous c-MYC-HSF1 interactions were detected by brightfield PLA in HeLa cells, following treatment with or without EtBr (100 µg/mL) for 1 hr. Scale bars: 10µm.

Demonstrating the specificity of PLA, *HSF1* KD markedly diminished the PLA signals. To validate direct c-MYC-HSF1 interactions *in vitro*, we performed Lumit immunoassays using recombinant proteins, where protein-protein interactions are indicated by the successful complementation of split NanoLuc® luciferase that are conjugated with two distinct antibodies^34^. Consistent with the co-IP and PLA results, GST-HSF1 did interact with c-MYC/MAX heterodimers *in vitro* compared to GST controls, evidenced by markedly elevated luminescence signals (Figure 3C). Next, we asked whether HSF1 can impact the interactions between c-MYC and MAX. Interestingly, HSF1 impaired the luciferase complementation denoting c-MYC-MAX interactions (Figure 3D). This finding suggests that HSF1 either induced conformational changes in the c-MYC/MAX heterodimer or simply blocked the recognition of c-MYC/MAX by antibodies. Nonetheless, either case supports a physical interaction between HSF1 and c-MYC/MAX dimers, which is further evidenced by *in vitro* pull-down assays. Recombinant His-HSF1 proteins were pulled down by GST-tagged c-MYC proteins, but not by GST proteins alone (Figure S3A). *Vice versa* was also true (Figure S3B). Moreover, these pull-down assays reveal that c-MYC alone can interact with HSF1.

Do HSF1 interactions affect the DNA binding of c-MYC/MAX dimers? To address this, we took advantage of a simple *in vitro* system, where recombinant c-MYC/MAX dimers can directly bind to DNA oligos containing the canonical E-box element that were immobilized on ELISA microtiter plates. This system was validated for capturing endogenous c-MYC/MAX dimers from nuclear extracts of MEFs with and without competition of free E-box elements (Figure S3C). Compared to GST controls, co-incubation with GST-HSF1 enhanced the binding of c-MYC/MAX dimers to E-box elements by over 60% (Figure 3E). This finding concurs with our cellular studies (Figures 1F and 1G).

### The c-MYC-MAX-HSF1 complex assembles on genomic DNAs

Whereas PLA can readily detect endogenous c-MYC-HSF1 interactions, co-IP of both has been technically challenging. Given the exclusive nuclear localization of PLA foci, we considered the possibility that the c-MYC/MAX-HSF1 complex might preferentially assemble on genomic DNAs. Therefore, regular cell lysis conditions would largely disrupt their associations.

First, we asked whether DNA binding is required for the interaction between HSF1 and c-MYC/MAX. To test this, we treated HEK293T cell lysates overexpressing FLAG-HSF1, HA-c-MYC, and V5-MAX with Ethidium bromide (EtBr). EtBr is known to disrupt DNA-dependent protein associations^35^. Of note, the whole cell lysates were prepared by sonication, under which genomic DNA fragments were present. EtBr treatment markedly abolished the interaction between HSF1 and c-MYC/MAX (Figure 3F), suggesting the dependency on genomic DNA binding. To exclude the possible contribution of cellular RNAs, RNase and DNase were applied to digest relevant substrates in these cell lysates, respectively. Treatment with DNase, but not RNase, disrupted the complex assembly (Figure 3G), demonstrating the necessity of genomic DNA binding. Of note, co-IP experiments cannot exclude the possibility that c-MYC and HSF1 may be brought together via their co-occupancy of adjacent genomic DNAs (Figure 3H).

However, this scenario would predict: 1) HSF1 DNA binding is required for c-MYC transcriptional activity; and 2) HSF1 and c-MYC lack physical interactions. Apparently, both predictions have already been refuted (Figure 1D and 3B). To further demonstrate the dependency on DNA binding at physiological conditions, bright field PLA was performed *in situ* to avert potential interference from EtBr fluorescence. The results confirmed a direct interaction between endogenous c-MYC and HSF1 in HeLa cells, which, importantly, was largely disrupted by EtBr treatment (Figure 3I). Collectively, these findings support nuclear assembly of c-MYC-MAX-HSF1 complexes, a physiological event markedly facilitated by genomic DNA binding.

### HSF1 activates c-MYC transcriptional activity via GCN5

How does HSF1 promote c-MYC DNA binding and transcriptional activation? Chromatin structure/topography affects the accessibility of genomic DNAs to transcription factors^11^. It was reported that c-MYC can recruit chromatin-modifying complexes, such as the STAGA co-activator complex containing the histone acetyltransferase GCN5, to remodel chromatin structures^8, 36^.

First, we asked whether GCN5 is important to c-MYC transcriptional activity by knocking down *Gcn5* in MEFs. Resembling *Hsf1* deficiency, *Gcn5* KD diminished the expression of c-MYC target genes (Figure 4A). A similar result was also obtained from the c-MYC reporter assay (Figure S4A), indicating that GCN5 is crucial to c-MYC transcriptional activity. Next, we asked whether HSF1 activates c-MYC via GCN5. As demonstrated above (Figure 1B), both the full-length HSF1^1–529^ and transcription-deficient HSF1^324–529^ mutants enhanced c-MYC activity; however, this activation was largely blocked by *GCN5* KD (Figure 4B), indicating a requirement for GCN5. Conversely, *GCN5* overexpression activated c-MYC without elevating its protein levels (Figure S4B). Of note, GCN5 overexpression was sufficient to rescue the diminished DNA binding of c-MYC in *Hsf1^CKO^* MEFs (Figure 4C).

**Figure 4:**
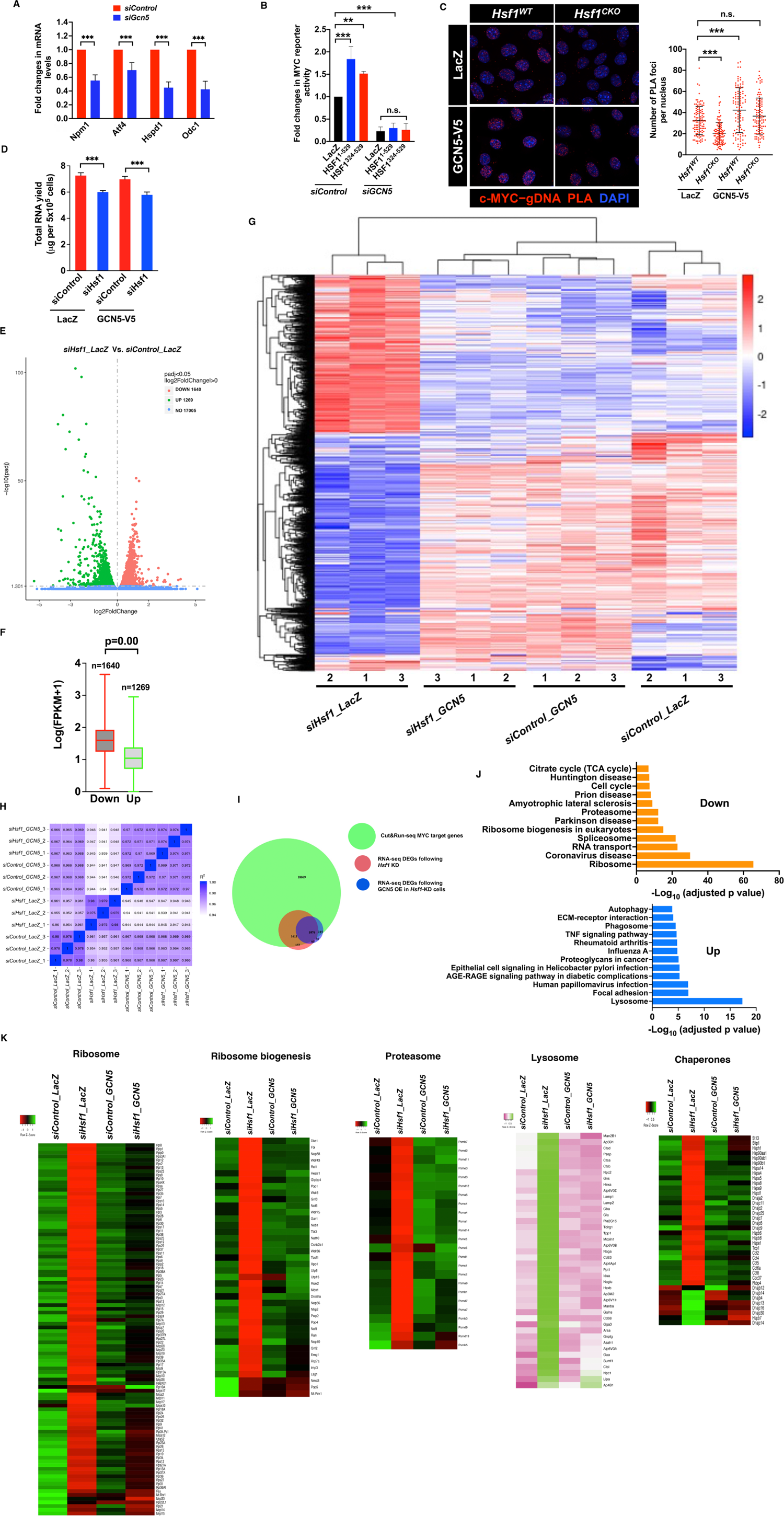
HSF1 activates c-MYC via GCN5. (A) The expression of known c-MYC target genes was quantitated by qRT-PCR, following transient *Gcn5* KD for 48 hr in immortalized MEFs (mean ± SD, n = 3 independent experiments, Two-way ANOVA). (B) Endogenous c-MYC transcriptional activities were measured by the dual reporter system in HEK293T cells transfected with indicated plasmids and siRNAs (mean ± SD, n = 3 independent experiments, One-way ANOVA). (C) Left panel: Endogenous c-MYC binding to gDNA binding was detected by PLA in immortalized MEFs stably expressing *LacZ* or *GCN5*. Scale bars, 10 µm. Right panel: quantitation of these PLA foci per nucleus (mean ± SD, n≥100 nuclei, One-way ANOVA). (D) Quantitation of total RNAs extracted with immortalized MEFs stably expressing LacZ or GCN5 (mean ± SD, n = 3 biological replicates, One-way ANOVA). (E) Volcano plot of the differentially expressed genes due to *Hsf1* KD. (F) Box-and-whisker plots of the abundance of DEGs in the control cells (n=1,640 or 1,269, Mann-Whitney U test). The box bounds the IQR divided by the median and the whiskers extend to the minimum and maximum values. (G) Visualization of DEGs in MEFs expressing different genes and siRNAs by clustering heatmaps (three biological replicates each group). (H) Seaborn correlation heatmap of gene expression among different experimental groups. (I) Venn diagram showing the overlaps among the c-MYC CUT&RUN-seq target genes, the DEGs following *Hsf1* KD, and the DEGs rescued by GCN5 overexpression in immortalized MEFs. (J) Pathway enrichment analyses of the DEGs in immortalized MEFs following *Hsf1* KD. (K) Heatmap visualization of the DEGs involved in the ribosome, proteasome, lysosome, and chaperone pathways (each data point represents the average of three biological replicates).

To determine how widespread the impacts of HSF1 on c-MYC transcriptional activity are, we conducted RNA-seq experiments. To avoid potential interference of 4-OHT with transcription^37^, we resorted to *Hsf1* KD in MEFs (Figure S4C). Interestingly, extraction of total RNAs from equal numbers of MEFs revealed that *Hsf1* KD resulted in a 18% reduction in RNA levels (Figure 4D). To account for this difference, we incorporated ERCC RNA spike-in controls during RNA extraction. Following appropriate normalization, RNA-seq data analyses revealed that total 2,909 genes were differentially expressed, both up-regulated and down-regulated, between the control and *Hsf1*-KD groups (Figure 4E and Table S4). In line with the overall reduction in total RNAs following *Hsf1* KD, those down-regulated genes displayed considerably higher abundance than those up-regulated genes (Figure 4F and Table S5). These changes in gene expression were illustrated by clustering heatmaps; interestingly, GCN5 overexpression markedly reversed these changes (Figure 4G and Table S6-S7). Congruently, the cells with both *Hsf1* KD and GCN5 overexpression were more closely correlated with the control cells than the *Hsf1*-KD cells, in terms of gene expression (Figure 4H). These findings highlight a key role of GCN5 in the HSF1-mediated transcription under non-stressed conditions. In line with its regulation of c-MYC, RNA-seq revealed that *Hsf1* KD altered the expression of an array of known c-MYC target genes, which was reversed by GCN5 overexpression (Figure S4D). Importantly, these RNA-seq findings were further validated by qRT-PCR (Figure S4E).

Next, we asked how many of these differentially expressed genes (DEGs) are c-MYC target genes. Our studies show that approximately 92% (2,687 out of 2,909) of those DEGs identified by RNA-seq are c-MYC target genes; moreover, GCN5 overexpression rescued the expression of nearly 40% of those 2,687 genes to varying degrees (Figure 4I), highlighting an important role of GCN5 in the specific regulation of c-MYC by HSF1.

Of interest, the differentially expressed c-MYC target genes following *Hsf1* KD play key roles in proteome homeostasis. Particularly, genes involved in the ribosome, ribosome biogenesis, proteasome and chaperone pathways are down-regulated; by contrast, genes involved in the lysosome and autophagy pathways, are up-regulated (Figures 4J and 4K). This gene up-regulation is not surprising, as c-MYC has been known to mediate transcriptional repression as well^38^. Whereas *Hsf1* KD altered the expression of chaperones that are constitutively expressed, these changes were reversed by GCN5 overexpression (Figure 4K), in line with a c-MYC-dependent mechanism. By contrast, c-MYC exhibited no or only low occupancy at the promoters of classic stress-inducible *Hsp* genes, including *Hspb1/Hsp25* and *Hspa1a/Hsp72* (Figure S4F). Compared to their constitutive cognates, their expression is either low or undetectable under non-stressed conditions (Figure S4F), as expected. Importantly, the diminished *Hspb1* expression, due to *Hsf1* KD, could not be rescued by GCN5 overexpression (Figure S4F), suggesting a c-MYC-independent, HSF1-dependent mechanism. In further support of our findings, approximately 74% of the DEGs following *Hsf1* KD in our MEFs are also differentially expressed in human medulloblastoma cells following *c-MYC* KD^39^ (Figure S4G). Collectively, these findings uncover a genome-wide impact of HSF1 on the c-MYC-mediated transcriptional program.

### HSF1 directly recruits GCN5 to c-MYC

Given the critical role of GCN5 in HSF1-mediated c-MYC regulation, we asked whether HSF1 influences the GCN5 recruitment to c-MYC. When overexpressed in HEK293T cells, FLAG-HSF1 was co-IPed with V5-GCN5 (Figure 5A). Although this finding suggests a direct recruitment of GCN5 by HSF1, it remains possible that HSF1 promotes c-MYC-GCN5 interactions indirectly. To distinguish these two possibilities, *in vitro* pull-down assays were performed using recombinant proteins. Compared to recombinant EHMT2 controls, a histone methyltransferase^40^, recombinant HSF1 proteins directly pulled down recombinant GCN5 proteins (Figure S5A), in support of a direct recruitment. This finding predicts that HSF1 deficiency would diminish the GCN5 association with c-MYC. Congruently, PLA indicated a reduced interaction between endogenous c-MYC and GCN5 in HeLa cells following *HSF1* KD (Figure 5B). Moreover, in MEFs *Hsf1* KD also impaired c-MYC-GCN5 association (Figure 5C). Conversely, HSF1 overexpression heightened their association (Figure S5B). Thus, these findings support a direct recruitment of GCN5 by HSF1 to c-MYC.

**Figure 5.**
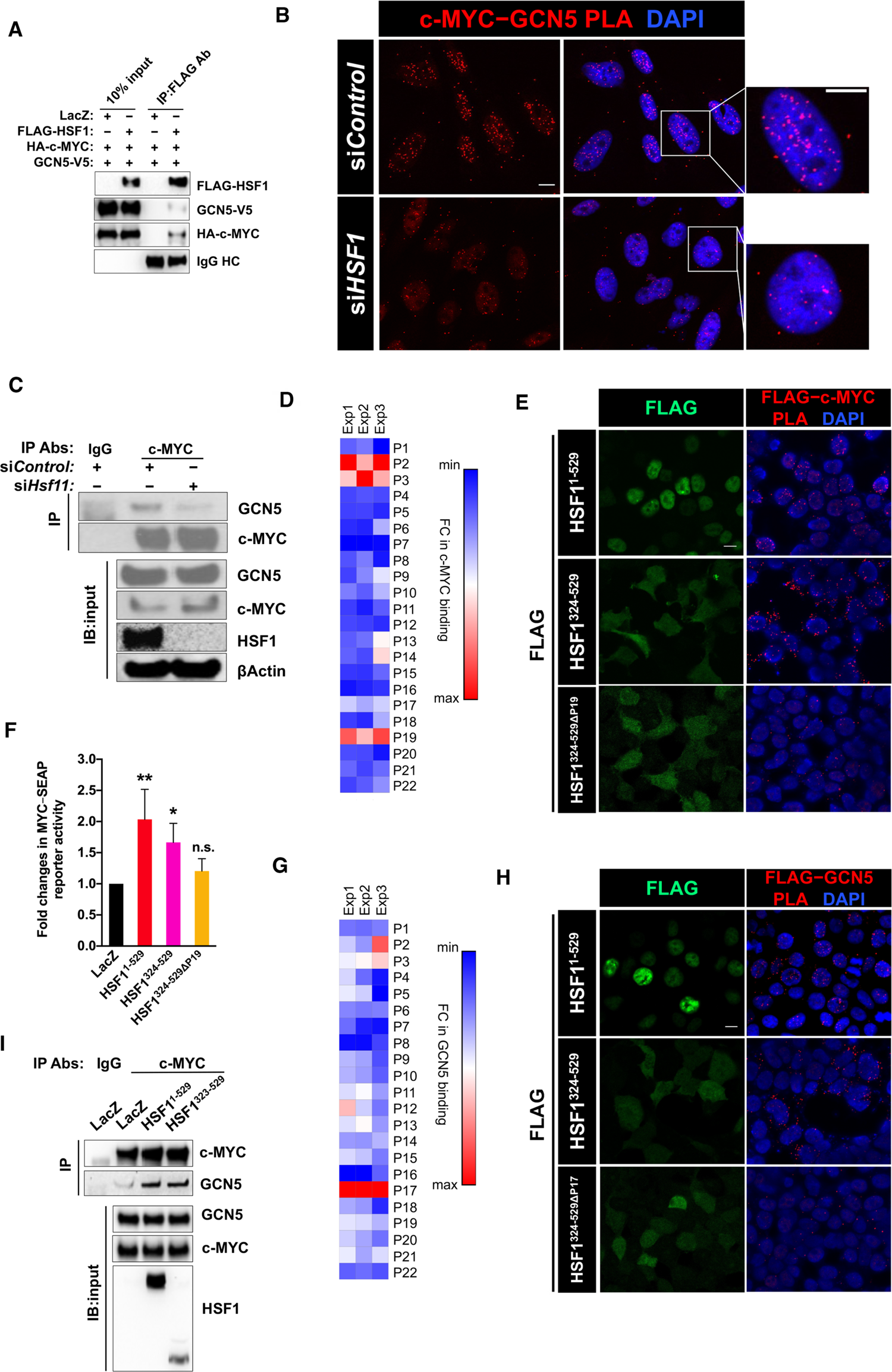
HSF1 recruits GCN5 to c-MYC. (A) Co-IP of FLAG-HSF1, HA-c-MYC, and V5-GCN5 in transfected HEK293T cells (representative images of three independent experiments). (B) Endogenous c-MYC-GCN5 interactions were detected by PLA in HeLa cells. Scale bars, 10 µm. (C) Co-IP of endogenous c-MYC and GCN5, following transient *Hsf1* KD in immortalized MEFs (representative images of three independent experiments). (D) *In vitro* binding of recombinant c-MYC proteins to individual HSF1 peptides immobilized on ELISA plates. Fold changes in binding are presented as a heatmap (n=3 independent experiments). (E) Visualization of interactions between transfected FLAG-HSF1 and endogenous c-MYC by PLA in HEK293T cells using a mouse monoclonal anti-FLAG antibody and a rabbit anti-c-MYC antibody. Scale bars, 10 µm. (F) c-MYC transcriptional activities were measured by the dual reporter system in HEK293T cells co-transfected with indicated plasmids (mean ± SD, n = 3 independent experiments, One-way ANOVA). (G) *In vitro* binding of recombinant GCN5 proteins to individual HSF1 peptides immobilized on ELISA plates. Fold changes in binding are presented as a heatmap (n=3 independent experiments). (H) Visualization of interactions between transfected FLAG-HSF1 and endogenous GCN5 by PLA in HEK293T cells. Scale bars, 10 µm. (I) Co-IP of endogenous c-MYC and GCN5 in HEK293T cells transfected with LacZ or FLAG-HSF1 (representative images of three independent experiments).

### HSF1 couples c-MYC and GCN5 via its C-terminal AD

Next, we embarked on elucidating the interactions among HSF1, c-MYC, and GCN5. To delineate the c-MYC binding sites on HSF1, we utilized a synthetic HSF1 peptide library, comprising 22 non-overlapping peptides (24 amino acids each), as described in our previous publication^20^. After screening for the binding of recombinant c-MYC proteins *in vitro*, three HSF1 peptides, located at the N-terminal DBD (P2, P3) and C-terminal AD (P19) respectively, displayed evident binding capability (Figure 5D). Considering that HSF1^1-3^^23^ was incapable of activating c-MYC (Figure 1D), we then focused on P19. Revealed by PLA, deletion of the P19 sequence largely abolished the interaction between FLAG-HSF1^324-529^ and endogenous c-MYC, supporting this region as the interacting interface with c-MYC (Figure 5E). Accompanied with this loss of interaction, P19 deletion abolished the HSF1-mediated c-MYC activation, indicating the necessity of their physical interaction (Figure 5F).

A similar screen was performed to delineate the GCN5 binding sites on HSF1. P17, another region located within the AD, was identified for strong GCN5 binding (Figure 5G). *In situ* PLA indicated that the P17 region was required for GCN5 binding, as its deletion markedly diminished FLAG-HSF1^324-529^-GCN5 interactions (Figure 5H). Importantly, overexpression of HSF1^324-529^, just like HSF1^1-529^, heightened the co-IP of c-MYC and GCN5 (Figure 5I). Together, our findings support that HSF1, via discrete interactions, couples GCN5 and c-MYC.

### HSF1 regulates the epigenetic state of c-MYC target loci

Chromatin remodeling is important to transcriptional regulation in eukaryotes. Given the diminished GCN5 recruitment to c-MYC, we predicted that histone acetylation mediated by GCN5 would be impaired in *Hsf1*-deficient cells. Consistently, ChIP experiments revealed that acetylation of H3K9/14, hallmarks of active gene promoters^41, 42^, was diminished in *Hsf1^CKO^* MEFs. Of note, this reduction occurred specifically at c-MYC target loci, but not at non-target loci (Figure 6A). In light of the importance of recruiting GCN5 to c-MYC, we further predicted that fusion of the HSF1 C-terminal AD, containing the GCN5 binding site, to c-MYC would generate a “superactive” c-MYC mutant. Interestingly, this HSF1-c-MYC fusion consistently resulted in markedly elevated protein expression, likely due to protein stabilization, compared to the c-MYC wildtype. To better compare their transcriptional activities, less amounts of this fusion plasmid were transfected into HEK293T cells (Figure 6B). Despite this decreased expression, the HSF1-c-MYC fusion still demonstrated markedly heightened transcriptional activity compared to the wildtype, as predicted (Figure 6B).

**Figure 6.**
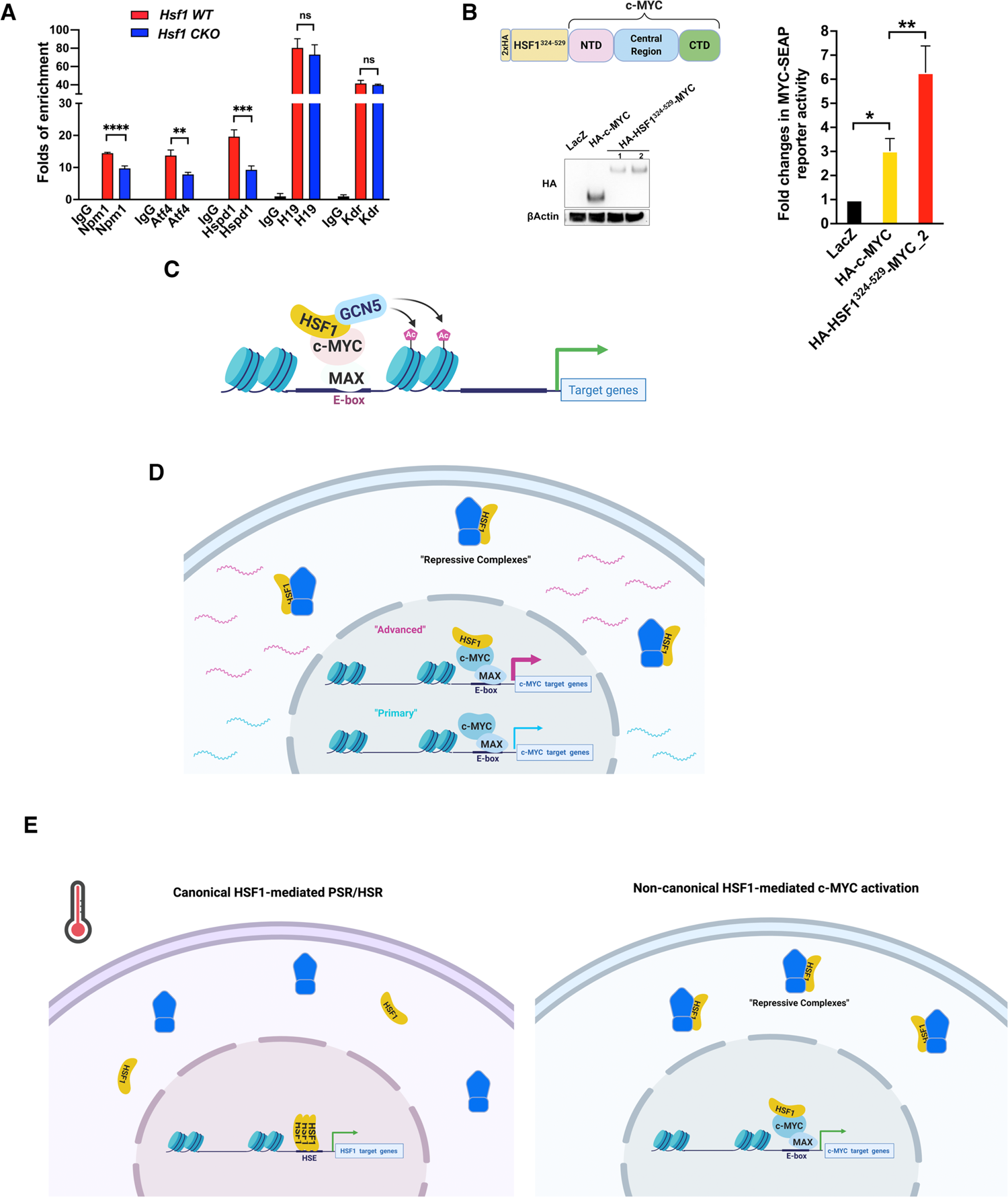
*Hsf1* deficiency impairs acetylation of histone H3 at c-MYC target loci. (A) ChIP-qPCR assays were performed to detect the acetyl-histone 3 (Lys9/Lys14) on c-MYC target or non-c-MYC target loci in immortalized MEFs (mean ± SD, n = 3 biological replicates, One-way ANOVA). (B) Left panel: the protein expression of fusion between HA-HSF1^324-529^ and c-MYC was detected by immunoblotting. Right panel: the transcriptional activity of fusion proteins was measured by the dual reporter system (mean ± SD, n = 3 independent experiments, One-way ANOVA). (C) The proposed model of HSF1-mediated c-MYC activation. HSF1 helps recruit GCN5 to c-MYC, thereby promoting chromatin remodeling and potentiating the c-MYC-mediated transcription. (D) HSF1 regulates two distinct activation states of c-MYC. Without HSF1 association, the transcriptional activity of cellular c-MYC remains low, sustaining at a primary state; by contrast, HSF1 association renders c-MYC highly active, transiting to an advanced state. (E) HSF1 governs at least two discrete transcriptional programs. Upon its activation, either in the face of environmental stress or within malignant cells, HSF1 initiates the canonical PSR/HSR, a mechanism of action depending on HSE binding. By contrast, in the absence of environmental stress most cellular HSF1 remains repressed; however, some HSF1 associates with c-MYC and potentiates its mediated transcription, a mechanism of action independent of HSE binding.

In aggregate, these findings support a molecular model, wherein HSF1, by directly recruiting GCN5 to c-MYC, promotes histone acetylation at the c-MYC target loci specifically, thereby heightening c-MYC DNA binding and, ultimately, magnifying its transcriptional activity (Figure 6C).

## DISCUSSION

Owing to its extensive regulation of the genome, potent oncogenic potential, and prominent role in pluripotency reprogramming, c-MYC has attracted great attention in biomedical research. Herein, we report that HSF1, a potent enabler of oncogenesis, specifically potentiates the c-MYC-mediated transcription. Our studies uncover a previously unrecognized transcription factor complex comprising both HSF1 and c-MYC/MAX heterodimers. Instead of binding to HSEs, unexpectedly, within this complex HSF1 directly recruits the histone acetyltransferase GCN5 to c-MYC via physical interactions. GCN5, in turn, remodels chromatin architecture to stimulate c-MYC transcriptional activity. Thereby, HSF1 renders c-MYC transcriptionally competent.

### A conditional, DNA binding-dependent transcription factor complex

Distinct from “constitutive” protein complexes, the assembly of c-MYC-MAX-HSF1 complexes is “conditional”, mainly contingent upon DNA binding. Although there is a possibility that HSF1 and c-MYC/MAX dimers may co-occupy adjacent genomic DNAs independently, several lines of evidence collectively refute this as a principal mechanism: 1) if this scenario were true, preventing HSF1 DNA binding would abolish their co-IP and c-MYC activation. Contrary to this prediction, HSF1^324–529^ mutants lacking the DBD are still able to co-IP with and activate c-MYC; 2) the PLA signals unequivocally denote a direct contact between HSF1 and c-MYC in intact cells; 3) importantly, the P19 region on HSF1 AD mediates c-MYC interactions; and 4) compared to E-boxes, HSEs were far less enriched in the c-MYC binding sites. Our data suggest that monomeric HSF1 is sufficient to associate with c-MYC/MAX, as HSF1^324-529^ mutants, which also lack the trimerization domain, still interacts with c-MYC. Nonetheless, we cannot exclude the possibility that trimeric HSF1 may also bind to c-MYC/MAX dimers especially under heat shock.

Furthermore, this conditional, DNA binding-dependent complex differs from the previously described “enhanceosome”^43^, where individual transcription factors cooperatively bind to their respective DNA elements. By contrast, while within this complex only c-MYC/MAX dictate the specificity of DNA binding, HSF1 behaves like an adaptor devoid of DNA binding. In a sense, this transcription factor complex operates in a “hybrid” mode, fusing the DNA binding capability of c-MYC/MAX with the transcription coregulatory function of HSF1. Owing to the conditional nature of this complex, HSF1 would not become limited for its *de facto* transcriptional program, namely the HSR/PSR, whilst amplifying the c-MYC-mediated transcription. Nonetheless, it remains elusive how this complex assembly depends on DNA binding. It is possible that DNA binding may incite conformational changes in c-MYC/MAX dimers, which, in turn, favors the interaction with HSF1. Further investigations are warranted. Unlike its dependency on DNA binding at the cellular context, this c-MYC-HSF1 interaction can be readily detected *in vitro* using recombinant proteins in the absence of DNA binding. This is most likely due to excessive proteins under *in vitro* conditions, bypassing the requirement for DNA binding. Under physiological conditions, however, cellular HSF1 and c-MYC proteins are either limited or unavailable for interaction, making DNA binding a prerequisite for efficient complex assembly.

It appears that at physiological conditions only part of cellular c-MYC/MAX dimers associate with HSF1. Of interest, the genomic loci of c-MYC targets regulated by HSF1 are enriched for histone acetylation, compared to non-HSF1-regulated targets (Figure S6A). Particularly, H3K27ac is a well-known epigenetic mark for active/open chromatins^44^. Consistent with preferential c-MYC/MAX DNA binding at these genomic loci, the CUT&RUN-seq binding sites display higher peak signals (Figure S6B). In support of active transcription, these HSF1-regulated c-MYC target genes are expressed at significantly higher levels (Figure S6C). To date, two distinct models of c-MYC-mediated transcription have been proposed: a gene selective activator (initiation) or a universal amplifier (elongation)^45^. Of note, our studies were conducted under physiological conditions without c-MYC overexpression. While our findings do not distinguish these two models, they collectively support a scenario wherein cellular c-MYC/MAX dimers preferentially bind to genomic loci possessing more open chromatin structures, which is ensued by the recruitment of HSF1 and GCN5 that stabilizes DNA binding and, ultimately, leads to enhanced transcriptional initiation or elongation. By forming this hybrid transcription factor complex, HSF1 not only empowers the c-MYC-mediated transcription but also greatly expands its own biological impacts, far beyond protein quality control.

### HSF1 dictates two distinct c-MYC activation states

Of interest, the ability of HSF1^324-529^ to directly recruit GCN5 may account for the effectiveness of HSF1 AD in newly emerged CRISPR activation systems^46^. Despite its necessity for the HSF1-mediated c-MYC activation, some GCN5 still associates with c-MYC even in the absence of *Hsf1*. Thus, HSF1 only augments the GCN5 association. This is crucial, considering that *c-MYC* is an essential gene. Therefore, *Hsf1*-deficient cells would retain a diminished c-MYC activity that is still sufficient to sustain viability. It remains to be determined whether c-MYC *per se* could recruit GCN5 independently. Conceptually, at the cellular level c-MYC activity could be retained at two distinct states, primary and advanced (Figure 6D). HSF1 controls the switch between these two. By engaging extra GCN5, HSF1 empowers c-MYC to function at its full capacity, which may be required for certain physiological and pathological conditions beyond simple viability maintenance.

HSF1 is dispensable for the viability of non-transformed cells, suggesting that the primary state of c-MYC activation is sufficient for viability. It further implies that the c-MYC-bound genes in *Hsf1^CKO^* cells may represent the core targets critical for life. In line with this notion, these 6,927 genes are enriched for common essential genes defined by Project Achilles and display higher probabilities of dependency in general (Figures S6D and S6E). Congruently, the gene ontology enrichment analysis reveal that these target genes engage in many essential biological processes, including ribosome biogenesis and mRNA processing (Figure S6F).

### HSF1 is a guardian of cellular proteome

It has been widely recognized that under stressed conditions HSF1 is crucial to the maintenance of proteomic stability through direct induction of *HSP* gene transcription. This action mainly protects protein quality. Now, our studies reveal that HSF1 can control protein quantity as well at both the synthesis and degradation phase. Through c-MYC, HSF1 transcriptionally regulates ribosomes, proteasomes, and lysosomes. Intriguingly, HSF1 governs not only translation capacity via ribosomes, indicated in this study, but also translation efficiency via mTORC1, as reported previously^31^.

Another interesting finding is the regulation of constitutively expressed *HSPs* by HSF1. Apart from its essential role in determining the expression of stress-inducible *Hsp* genes, namely the HSR/PSR, HSF1 also augments the expression of constitutively expressed *Hsp* genes via c-MYC. Thus, by overseeing every major aspect of proteome homeostasis, HSF1 acts as a guardian of cellular proteome.

### Implications in stress, cancer, and stem cell biology

Canonically, the HSR/PSR is characterized by the specific binding of HSF1 trimers to HSEs located at gene promoters and subsequent transcriptional induction of these target genes, many of which encodes HSPs. Although HSF1 can regulate *non-HSP* genes, including the target genes of E2F^47, 48^, this regulation is also reliant on the HSE binding of HSF1. Apparently, this HSF1-c-MYC complex does not follow this classic definition. Independently of DNA binding, HSF1 can activate the much broader c-MYC-mediated transcriptional program (Figure 6E), exerting more profound impacts on cellular physiology than previously thought. Of note, under non-stressed conditions most HSF1 remains repressed and inactive; however, some HSF1 appears to escape this repression and potentiate the c-MYC-mediated transcription, independently of HSE binding (Figure 6E). Thus, even in the absence of proteotoxic stress HSF1 remains transcriptionally active to impact cellular physiology. Moreover, the versatility of HSF1 to direct distinct transcriptional programs, depending on different complexes it forms, exemplifies a new mode of action of transcription factors.

Ample evidence has pinpointed HSF1 as a generic pro-oncogenic factor, via multifaceted mechanisms^16–21^. Of note, in non-transformed MEFs *Hsf1* deficiency affected the expression of roughly 12% of the c-MYC target genes, although it is likely underestimated due to incomplete *Hsf1* KD. This finding suggests that only part of cellular c-MYC is associated with HSF1 under this condition. Likely, in non-transformed cells HSF1 is largely inaccessible, partly due to its repressive mechanisms, to activate c-MYC. However, in human cancers HSF1 is notably overexpressed^14, 49^. This increased quantity would render a considerable portion of cellular c-MYC transcriptionally competent, thereby promoting malignancy. In support of this notion, approximately 80% of HSF1-bound genes, defined by HSF1 ChIP-seq^49^, in human cancers are c-MYC targets (Figure S6G). Given that in cancerous cells HSF1 becomes constitutively active^14, 50^, the rest 20% likely comprise canonical HSF1 targets. Conversely, without HSF1, cells only possess basic c-MYC activity that is sufficient for viability but inadequate for malignant transformation, thus adopting a “tumor-resistant” cellular state. This concept may have implications in anti-cancer therapies. Owing to its essentiality to viability, directly targeting c-MYC likely inflicts undesirable side effects. Instead, targeting HSF1 may abate c-MYC activity to a level that is adequate to sustain viability, but unable to support malignancy. Excitingly, novel HSF1 inhibitors showing potent anti-cancer effects have been developed in recent years^51, 52^.

Lastly, given the importance of c-MYC to pluripotency reprogramming, it is plausible to postulate that this HSF1-mediated c-MYC activation may impact stemness. Although HSF1 has been implicated in maintaining cancer stem cells^53, 54^, its role in normal stem cell biology remains to be determined.

## EXPERIMENTAL MATERIALS AND METHODS

### Cell culture and reagents

HeLa cells were purchased from ATCC and HEK293T cells were purchased from GE Dharmacon. Both were recently authenticated by ATCC. Immortalized *Rosa26-CreER^T2^; Hsf1^fl/fl^* MEFs (male) were described previously^31^. To delete *Hsf1*, these MEFs were pre-treated with ethanol or 1 mM (Z)-4-Hydroxytamoxifen (4-OHT) for 7 days. A2058 cells stably expressing LacZ or FLAG-HSF1 were described previously^16^. All cell cultures were maintained in DMEM supplemented with 10% HyClone bovine growth serum and 1% penicillin–streptomycin (Gibco). Cells were maintained in an incubator with 5% CO2 at 37 °C. All cell lines were routinely tested for mycoplasma contamination using MycoAlert Mycoplasm Detection kits.

Recombinant proteins were all purchased commercially, including c-MYC/MAX complexes (Cat#81087, Active Motif Inc.), GST (Cat#G52-30U, SignalChem Biotech), GST-HSF1 (Cat#H25-30G, SignalChem Biotech), His-HSF1 (Cat#ADI-SPP-900, Enzo Life Sciences Inc.), GST-c-MYC (Cat#H00004609-P01, Abnova Corp.), His-c-MYC (Cat#230-00580-100, RayBiotech, Inc.), His-GST (Cat#12-523, Sigma-Aldrich Inc.), FLAG-EHMT2 (Cat#31410, Active Motif Inc.), and FLAG-GCN5 (Cat#31591, Active Motif Inc.).

### Plasmids and generated stable cells

pBabe-HSF1-FLAG was a gift from Robert Kingston (Addgene plasmid#1948). pMSCV-HA-cMYCT58A was a gift from Scott Lowe (Addgene plasmid#18773). pCherry-HSP90alpha was a gift from Didier Picard (Addgene plasmid#108222). pCDNA3-2xHA-c-MYC was a gift from Martine Roussel (Addgene plasmid#74161). pLX304-LacZ-V5 was a gift from William Hahn (Addgene plasmid#42560). pBabe-LacZ, pBabe-HSF1^1-323^, and pBabe-HSF1^324-529^ were described previously^20^. pLX304-MAX-V5 (HsCD00440967) and pDONR221-GCN5 (HsCD00829789) vectors were purchased from DNASU plasmid depository. pLX304-LacZ-V5 and pLX304-GCN5-V5 vectors were co-transfected with packaging vector (delta VPR) and an envelope vector (VSV-G) into HEK293T packaging cells using TurboFect transfection reagent (Cat#R0531, ThermoFisher). MEF cells were infected with produced lentivirus in the presence of polybrene (10 µg/mL). After incubation for 3 days, cells were selected with 1 µg/mL blasticidin for 7 days.

### Transfection and c-MYC dual reporter assays

All plasmids were transfected with TurboFect transfection reagents. HEK293T cells were co-transfected with pMYC-SEAP and pCMV-Gaussia luciferase (GLuc) reporter plasmids, along with various indicated plasmids. After 48 hours, reporter activities in culture media were measured. SEAP and GLuc activities in culture supernatants were quantitated using a NovaBright Phospha-Light EXP Assay Kit (Cat#N10577, ThermoFisher Scientific) for SEAP and a Pierce™ Gaussia Luciferase Glow Assay Kit (Cat#16160, ThermoFisher Scientific), respectively. Luminescence signals were measured by a CLARIOstar microplate reader (BMG LABTECH). SEAP activities were normalized against GLuc activities.

### siRNA and shRNA knockdown

siRNAs were transfected at 10nM final concentration, except *c-Myc*-targeting siRNAs (50 nM final concentration), using Mission® siRNA transfection reagent or jetPRIME*®* transfection reagent. siRNAs were purchased commercially, including non-targeting control siRNAs (Cat#D-001210-02-05, Horizon Discovery Ltd.), *Hsf1*-targeting siRNAs (Cat#SASI_Mm01_00023056 and _00023057, Signa-Aldrich), *HSF1*-targeting siRNAs (Cat# SASI_Hs01_00067735 and _Hs02_00339745, Signa-Aldrich)*, c-Myc*-targeting siRNAs (SASI_Mm01_00157474 and _00157475, Signa-Aldrich), *Gcn5*-targeting siRNAs (Cat# SASI_Mm01_00159517 and Mm02_00289578, Signa-Aldrich), and GCN5-targeting siRNAs (Cat# SASI_Hs01_00050954 and _00050955, Signa-Aldrich).

### Quantitative real-time PCR

Total RNAs were isolated using RNA STAT-60™ reagent (Cat#CS110, Tel Test Inc.), and 1 µg RNAs were used for reverse transcription using iScript™ cDNA Synthesis Kit (Cat#1708891, Bio-Rad). Equal amounts of cDNA were used for quantitative RCR reaction using a DyNAmo SYBR Green qPCR kit (Cat#F410L, ThermoFisher Scientific). Signals were detected by an Agilent Mx3000P qPCR System (Agilent Genomics). ACTB was used as the internal control. The sequences of individual primers for each gene are listed in Table S8.

### Immunoblotting and Immunoprecipitation

Whole-cell protein extracts were prepared in cold cell-lysis buffer (100 mM NaCl, 30 mM Tris-HCl pH 7.6, 1% Triton X-100, 20 mM sodium fluoride, 1mM EDTA, 1mM sodium orthovanadate, and 1x Halt™ protease inhibitor cocktail). Proteins were transferred to nitrocellulose membranes. Following incubation with the blocking buffer (5% non-fat milk in 1x TBS-T) for 1 hour at RT, membranes were incubated with primary antibodies (1:1,000 dilution in the blocking buffer) overnight at 4 °C. After washing with 1xTBS-T for 3 times, membranes were incubated with peroxidase-conjugated secondary antibodies (1: 5000 dilution in the blocking buffer) at RT for 1 hr. Signals were detected using SuperSignal West chemiluminescent substrates (Cat#34578 or #34095, ThermoFisher Scientific). For Co-IP, 1 mg whole cell lysates were incubated with primary antibodies at 4 °C overnight. Either normal rabbit IgG were used as the negative controls. Protein G magnetic beads (Cat#88847, ThermoFisher Scientific) were used to precipitate primary Abs. After washing with the lysis buffer for 3 times, beads were boiled in 1x loading buffer for 5 min before loading on SDS-PAGE.

### *In vitro* Lumit™ Immunoassays

The storage buffers of recombinant proteins were first changed to 1x Lumit™ Immunoassay buffer C using Zeba™ Spin desalting columns (7K MWCO, Cat#89883, ThermoFisher Scientific Inc.). For each reaction, 10 ng recombinant c-MYC/MAX complexes (Cat#81087, Active Motif Inc.) were incubated at 1:1 molar ratio with either recombinant GST (Cat#G52-30U, SignalChem Inc.) or GST-HSF1 proteins (Cat#H25-30G, SignalChem Inc) in 50 μl 1x Lumit™ Immunoassay buffer C at RT for 1 hr with 200rpm shaking. Τhen, 50 μl 1x Lumit™ Immunoassay buffer C containing 150 ng primary antibodies, including a rabbit anti-FLAG antibody (Cat#14793S, Cell Signaling Technology) in combination with a mouse anti-GST (26H1) antibody (Cat# 2624S, Cell Signaling Technology) for c-MYC-HSF1 interactions, or a mouse anti-FLAG (9A3) antibody (Cat#8146S, Cell Signaling Technology) in combination with a rabbit anti-His tag (D3I1O) antibody (Cat#12698S, Cell Signaling Technology) for c-MYC-MAX interactions, and 150 ng Lumit™ secondary antibodies was added to each well and incubated at RT for 90 min. Following the incubation, 25 μl 1x Lumit™ Immunoassay buffer C containing Lumit™ substrate C (1:12.5 dilution) in was added to each well and incubated for 2 min with 400 rpm shaking. The luminescence signals were measured by a SpectraMax iD5 microplate reader (Molecular Device, Inc.).

### c-MYC DNA binding assay

c-MYC DNA binding was measured by TransAM™ c-MYC transcription factor assay kits (Cat# 43396, Active Motif). The nuclei of MEFs were prepared by NE-PER™ Nuclear and Cytoplasmic extraction reagents (Cat#78835, ThermoFisher Scientific, Inc.). Isolated nuclei were lyzed in the complete lysis buffer to extract nuclear proteins. Each well was incubated with 50 μg nuclear extracts with and without the competition of wild-type E-box oligonucleotides. The detection of DNA-bound c-MYC followed the manufacturer’s instructions.

The microplates from the TransAM™ c-MYC transcription factor assay kit, on which consensus E-box oligonucleotides have been immobilized, were adapted to measure the DNA binding of recombinant c-MYC/MAX proteins. First, 10 ng recombinant c-MYC/MAX complexes were incubated with either recombinant GST or GST-HSF1 proteins (1:1 molar ratio) in 50 μl 1x DNA binding buffer (10 mM Tris, 50 mM KCl, pH 7.5) at RT for 1 hr with rotation. Following the incubation, the mixtures were loaded on the microplates and incubated at RT for 30 min with 200rpm shaking. Then, 50 μl 1x DNA binding buffer containing anti-FLAG antibody HRP conjugates (1:1000 dilution) was added to each well and incubated at RT for 15 min with 200rpm shaking. After 5 times of washing with 1x DNA binding buffer, 100 μl 1-Step Ultra TMB-ELISA Substrate Solution (Cat#34029, ThermoFisher Scientific Inc.) was added to each well for signal development.

### *In vitro* recombinant protein pull-down assay

400ng recombinant His-HSF1 (Cat#ADI-SPP-900, Enzo Life Sciences Inc.), FLAG-GCN5 (Cat#31591, Active Motif), FLAG-EHMT2 (Cat#31410, Active Motif), GST-MYC (Cat#H00004609-P01, Abnova) or His-GST (Cat#12-523, Millipore Sigma) were diluted in 400 µL reaction buffer (25mM Tris-HCL 100mM NaCl, 0.5% Triton X-100, pH7.5), followed by incubation for 3 hours at 4 °C. For the GST pulldown, glutathione magnetic beads (Cat#78601, ThermoFisher Scientific) were added and incubated at RT for 2 hours. For the other pulldowns, either rabbit anti-HSF1 (H-311) (Cat#sc-9144, Santa Cruz Biotechnology) or rabbit anti-FLAG antibodies (Cat#14793S, Cell Signaling Technology) were added to the mixtures and incubated for 3 hours at 4 °C, followed by incubation with protein G magnetic beads for 2 hours at 4 °C. Magnetic beads were collected and washed with reaction buffer, followed by protein elution (boiled in 1x sample buffer) and western blotting.

### Proximity Ligation Assay

Cells were fixed with 4% formaldehyde in PBS for 15 min at RT. After blocking with 5% goat or horse serum in PBS with 0.3% Triton X-100, cells were incubated with a pair of indicated rabbit, mouse, or goat primary antibodies (1:100 diluted in the blocking buffer) overnight at 4 °C. Following incubation with Duolink™ PLA anti-rabbit Plus, anti-mouse Minus, or anti-goat Minus probes (Cat#DUO92002, DUO92004, and DUO92006, Sigma-Aldrich) at 37 °C for 1 hour, ligation, rolling circle amplification, and detection were performed using Duolink™ In Situ Detection Reagents Red (Cat#DUO92008, Sigma-Aldrich). Nuclei were stained with Hoechst 33342. Signals were visualized using a Zeiss LSM780 confocal microscope. For brightfield PLA, detection was performed using Duolink™ In Situ Detection Reagents Brightfield (Cat#DUO92012, Sigma-Aldrich).

For the c-MYC-gDNA PLA, a rabbit anti-c-MYC (D3N8F) antibody (Cat#13987S, Cell Signaling Technology) was combined with a mouse anti-dsDNA (HYB331-01) antibody (Cat#sc-58749, Santa Cruz Biotechnology). For the c-MYC-HSF1 PLA, a rabbit anti-c-MYC (D3N8F) antibody (Cat#13987S, Cell Signaling Technology) was combined with a mouse anti-HSF1 (E-4) antibody (Cat#sc-17757, Santa Cruz Biotechnology). For the c-MYC-GCN5 PLA, a goat anti-c-MYC antibody (Cat#AF3696, R&D Systems) was combined with a rabbit anti-GCN5 (C26A10) antibody (Cat#3305S, Cell Signaling Technology). For the FLAG-HSF1-c-MYC PLA, a mouse anti-FLAG (9A3) antibody (Cat#8146S, Cell Signaling Technology) was combined with a rabbit anti-c-MYC (D3N8F) antibody (Cat#13987S, Cell Signaling Technology). For the FLAG-HSF1-GCN5 PLA, a rabbit anti-GCN5 (C26A10) antibody (Cat#3305S, Cell Signaling Technology) was combined with a mouse a mouse anti-FLAG (9A3) antibody (Cat#8146S, Cell Signaling Technology).

### Chromatin immunoprecipitation assay

The ChIP assay was performed using a SimpleChIP® Enzymatic Chromatin IP Kit (Cat#9003, Cell Signaling Technology) following the manufacturer’s instruction. Briefly, ∼4×10^6^ cells were fixed with 1% formaldehyde and quenched in glycine. Cells were lysed in extraction buffer to obtain nuclear pellet, followed by incubation with micrococcal nuclease to fragment genomic DNAs. Further sonication is performed to completely lyse the nuclei. Sheared DNAs were immunoprecipitated by normal rabbit IgG (Cat#10500C, ThermoFisher Scientific), rabbit c-MYC (D3N8F) monoclonal Abs (Cat#13987, Cell Signaling Technology), or rabbit Acetyl-Histone H3(Lys9/Lys14) Abs (Cat#9677, Cell Signaling Technology), followed by quantitative real-time PCR analysis. The total genomic DNAs immunoprecipitated by c-MYC Abs were measured using a DNA quantification fluorometric kit (Cat#K539, BioVision), following the manufacturer’s instruction. The sequences of oligos used for ChIP-qPCR are listed in Table S8.

### Detection of MYC/GCN5 binding by ELISA

The HSF1 peptide library was synthesized by GenScript Custom Peptide Synthesis Service. The amino acid sequences of individual peptides are listed in our previous publication^20^. Peptides were dissolved in 0.01N NaOH to make 1mM stocks. For detection of c-MYC/GCN5 binding sites, 20 mM HSF1 peptides in 100 µL PBS were coated on an ELISA microplate at 4 °C overnight. The plates were blocked with 1%BSA in PBS at RT for 30 min, followed by incubation with 20 ng recombinant c-MYC/MAX complexes or GCN5 proteins in 100 µL PBS-T buffer per well at 4 °C overnight. After washing with PBST for 3 times, each well was incubated with Rabbit anti-c-MYC (D3N8F) monoclonal Abs (Cat#13987, Cell Signaling Technology) or Rabbit anti-GCN5 monoclonal Abs (Cat#3305, Cell Signaling Technology) (1:1000 diluted in the blocking buffer) at RT for 3 hours. Following washing, each well was incubated with anti-Rabbit IgG (H+L)-HRP conjugates (1:5000 diluted in the blocking buffer) at RT for 1 hour. Signals were developed using the 1-Step Ultra TMB-ELISA Substrate Solution.

### RNA-seq and data analysis

MEFs stably expressing LacZ or V5-GCN5 were transfected with control or *Hsf1*-targeting siRNAs for 2 days. Total RNAs were extracted from 5×10^5^ MEFs, triplicates each experimental group, using Direct-zol RNA miniprep plus kit (Cat#R2073, Zymo Research). 1.5 μl of ERCC ExFold RNA spike-in mix 1 (1: 10 dilution, Cat#4456739, ThermoFisher Scientific Inc.) was added to each siControl RNA sample and 1.5 μl of mix 2 (1:10 dilution) was added to each *siHsf1* RNA sample. Libraries were prepared with rRNA depletion and sequenced with an Illumina HiSeq PE150 platform. Filtered raw data were mapped to the reference genome using HISAT2^55^. RUVseq package was used to normalize the data^56^. DESeq2 was used to analyze the DEG of samples^57^ (padj<=0.05 |log2FoldChange|>=0.0 are set as threshold). Hierarchical clustering was performed using the FPKMs of transcripts. Pathway enrichment analyses were performed using Enrichr^58^.

### CUT&RUN-seq and ChIP-seq

Cut&Run experiments were performed using a CUTANA™ ChIC/CUT&RUN kit (Cat# 14-1048, EpiCypher) according to the manufacturer’s instructions. Briefly, proliferating MEFs were crosslinked with 1% formaldehyde in PBS for 1 min on culture plates. After quenching with glycine, cells were scraped off the plates and counted. 5×10^5^ crosslinked cells were used for each sample. For the IgG control, both *Hsf1^WT^* and *Hsf1^CKO^* MEFs were mixed at a 1:1 ratio and incubated with rabbit IgG negative control antibodies. For the experimental groups, either *Hsf1^WT^* or *Hsf1^CKO^* MEFs (two biological replicates each group) were incubated with rabbit anti-c-MYC (D3N8F) monoclonal Abs (Cat#13987, Cell Signaling Technology). Of note, wash, cell permeabilization, and antibody buffers were all supplemented with 1% Triton X-100 and 0.05% SDS. Reversing cross-links was achieved by adding 0.8 μl of 10% SDS and 1 μl of 20 μg/μl Proteinase K to each sample and incubated at 55^0^C overnight. Following purification, 0.5 ng *E. coli* spike-in DNAs were added to each eluted DNA sample. Total 10 ng DNAs each sample were used to generate sequencing libraries using a NEBNext® Ultra™ II DNA Library Prep Kit for Illumina (Cat#E7645, New England Biolabs Inc.). The clustering of indexed samples was performed using a TruSeq PE Cluster kit v3-cBot-HS (Cat#PE-401-3001, Illumina, Inc.). The library preparations were sequenced on an Illumina HiSeq 2500 system to generate 150 bp paired-end reads. The sequencing data were analyzed using the EpiCypher Cut&Run pipeline (Basepair Inc.). Briefly, following trimming, the raw sequencing reads were aligned to the mouse (GRCm38/mm10) and *E. coli* (strain K-12) reference genomes respectively using Bowtie2^59^.

Subsequently, CUT&RUN peaks were called using SEACR^60^ with the stringent and spike-in normalization settings. As a comparison, CUT&RUN peaks were also called using MACS2^61^, which results in much fewer peaks (5075 for WT and 2198 for CKO). The motif enrichment analyses were performed using AME^62^.

The ChIP-seq experiments and data analyses were done through a contract with the Active Motif Epigenetic Services (Active Motif, Inc.). Briefly, equal amounts of sonicated chromatins from two biological replicates were used for ChIP using the same anti-c-MYC antibodies. Input chromatins were sequenced as the control. Paired-end reads were aligned to the mouse (GRCm38/mm10) reference genome using Bowtie2 and ChIP-seq peaks were called using MACS2.

### Statistical analyses

Statistical analyses were performed using Prism GraphPad 8.0 (GraphPad Software). The statistical significance is defined as: *p<0.05, **p<0.01; ***p<0.001; n.s.: not significant. For *in vitro* experiments, sample size required was not determined a priori. The experiments were not randomized.

## ACKNOWLEDGMENTS

We would like to thank the Optical Microscopy and Image Analysis lab (OMAL) for their assistance with the confocal microscopy studies. This work was supported by the grant from National Institutes of Health (NIH), United States to C.D. (1DP2OD007070) and by the Intramural Research Program of the NIH, National Cancer Institute, Center for Cancer Research. The content of this publication does not necessarily reflect the views or policies of the Department of Health and Human Services, nor does mention of trade names, commercial products, or organizations imply endorsement by the U.S. government.

## Author contributions

M. X., L. L., K-H. C., B. R., S. D., K-H. S., Z. T., and C. D. designed and conducted the experiments. C.D. conceptualized and supervised this study and analyzed the results. M.X. and C.D. wrote the manuscript.

