## Supplemental Figures 1-6 for "Heat Shock Factor 1 (HSF1) specifically potentiates c-MYC-mediated transcription independently of the canonical heat-shock response"

**LIST OF SUPPLEMENTARY MATERIALS**

The supplementary materials contain 6 supplementary Figures and their legends, and 8 supplementary tables.

**Figure S1: HSF1 is required for robust c-MYC-mediated transcription.**

**Figure S2: HSF1 promotes c-MYC DNA binding.**

**Figure S3: HSF1 physically interacts with c-MYC.**

**Figure S4: HSF1 activates c-MYC via GCN5.**

**Figure S5: HSF1 recruits GCN5 to c-MYC.**

**Figure S6: HSF1 potentiates the c-MYC mediated transcription.**

**Table S1: CUT&RUN-seq peaks called by SEACR in *Hsf1*$^{WT}$ MEFs.**

**Table S2: CUT&RUN-seq peaks called by SEACR in *Hsf1*$^{CKO}$ MEFs.**

**Table S3: ChIP-seq peaks called by MACS2 in *Hsf1*$^{WT}$ MEFs.**

**Table S4: RNA-seq DEGs in MEFs following *Hsf1* KD.**

**Table S5: RNA-seq gene expression (FPKM) of all four experimental groups.**

**Table S6: RNA-seq differential gene expression (FPKM) between *siHsf1_LacZ* and *siControl_GCN5* groups.**

**Table S7: RNA-seq differential gene expression (FPKM) between *siHSF1_LacZ* and *siHsf1_GCN5* groups.**

**Table S8: Nucleotide sequences of primers.**

### Figure S1

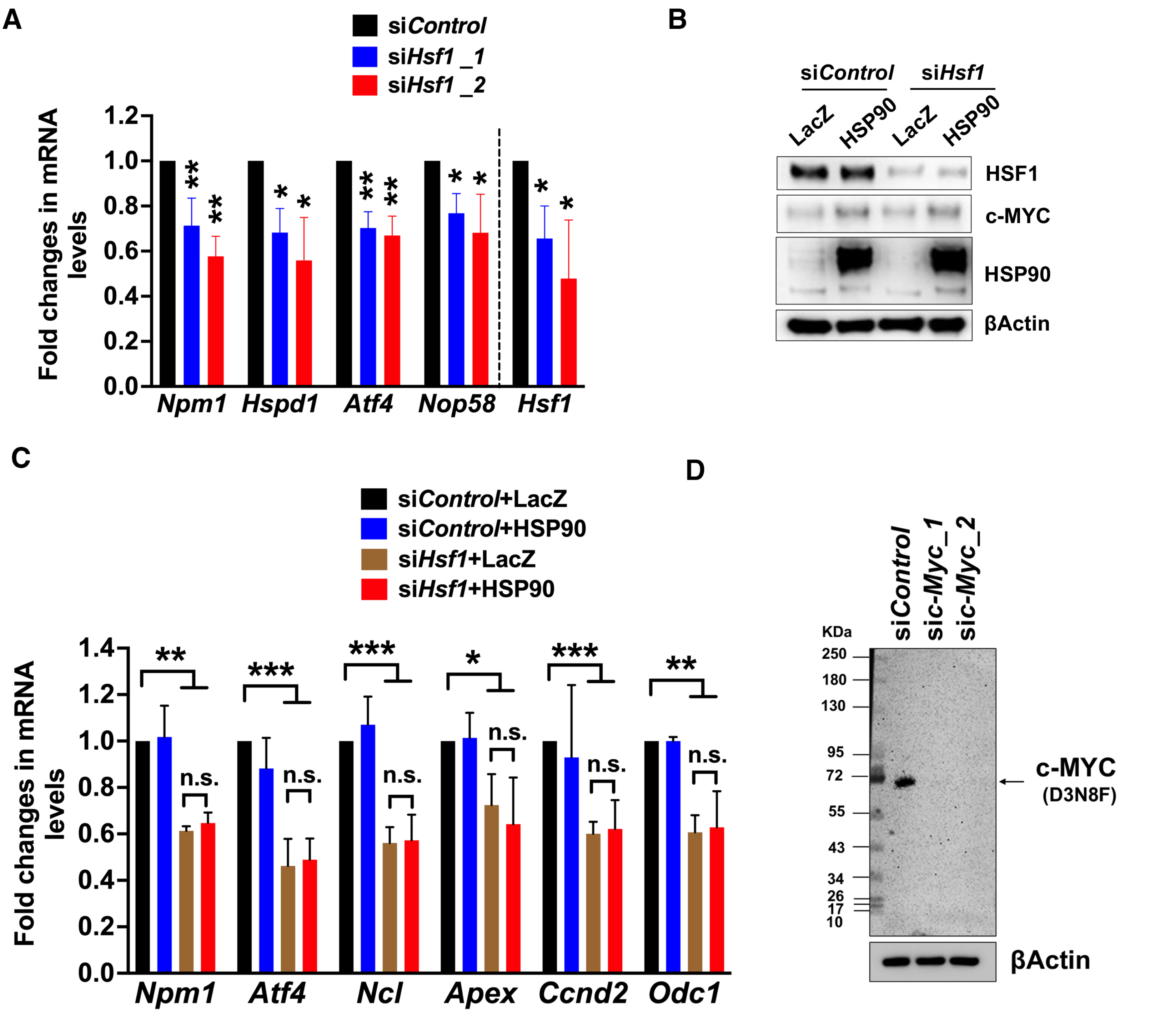
A
siControl
siHsf1 _1
siHsf1 _2
Fold changes in mRNA levels
1.2
1.0
0.8
0.6
0.4
0.2
0.0
Npm1
Hspd1
Atf4
Nop58
Hsf1
B
siControl
siHsf1
LacZ
HSP90
LacZ
HSP90
HSF1
c-MYC
HSP90
βActin
C
siControl+LacZ
siControl+HSP90
siHsf1+LacZ
siHsf1+HSP90
Fold changes in mRNA levels
1.4
1.2
1.0
0.8
0.6
0.4
0.2
0.0
**
***
***
*
***
**
n.s.
n.s.
n.s.
n.s.
n.s.
n.s.
Npm1
Atf4
Ncl
Apex
Ccnd2
Odc1
D
KDa
siControl
sic-Myc_1
sic-Myc_2
250
180
130
95
72
55
43
34
26
17
10
c-MYC
(D3N8F)
βActin

#### SUPPLEMENTARY FIGURE LEGENDS

**Figure S1**: **HSF1 is required for robust c-MYC-mediated transcription.**

(A) mRNA levels of known c-MYC target genes were quantitated by qRT-PCR, following transient knockdown of *Hsf1* in immortalized MEFs for 48 hr (mean ± SD, n =3 independent experiments, One-way ANOVA). (B) and (C) Knocking down *Hsf1* and overexpressing HSP90α in immortalized MEFs. Protein expression was detected by immunoblotting (B). mRNA levels of c-MYC target genes were quantitated by qRT-PCR (C) (mean ± SD, n =3 independent experiments, Two-way ANOVA). (D) Validation of the rabbit anti-c-MYC antibody (D3N8F). c-MYC protein levels were detected by immunoblotting in immortalized MEFs transfected with two independent siRNAs targeting *c-Myc* for 4 days.

### Figure S2

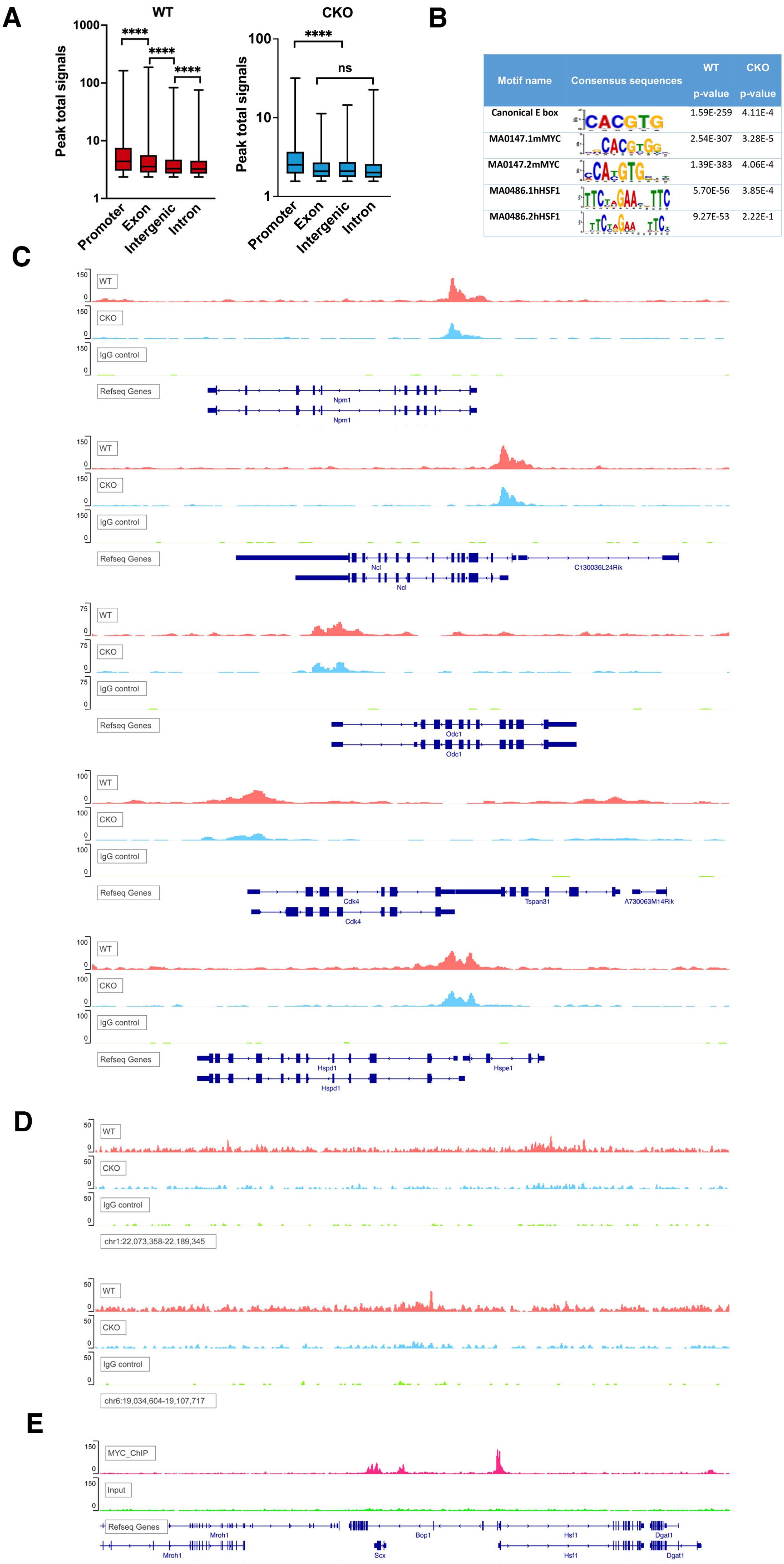

A
WT
CKO
Peak total signals
1000
100
10
1
****
****
****
ns
Promoter
Exon
Intergenic
Intron
B
Motif name
Consensus sequences
WT p-value
CKO p-value
Canonical E box CACGTG 1.59E-259 4.11E-4
MA0147.1mMYC 2.54E-307 3.28E-5
MA0147.2mMYC 1.39E-383 4.06E-4
MA0486.1hHSF1 5.70E-56 3.85E-4
MA0486.2hHSF1 9.27E-53 2.22E-1
C
WT
CKO
IgG control
Refseq Genes
Npm1
Ncl
C130036L24Rik
Odc1
Cdk4
Tspan31
A730063M14Rik
Hspd1
Hspe1
D
chr1:22,073,358-22,189,345
chr6:19,034,604-19,107,717
E
MYC_ChIP
Input
Mroh1
Bop1
Hsf1
Dgat1
Scx

**Figure S2: HSF1 promotes c-MYC DNA binding.**

(A) Box plots of peak signals based on their genomic locations. The box bounds the IQR divided by the median and the whiskers extend to the minimum and maximum values (n=18859, 24455, 110779, and 55187 peaks for WT, n=4090, 223, 968, and 600 peaks for CKO, One-way ANOVA). (B) Enrichment of c-MYC- and HSF1-binding motifs in peak sequences. (C) Visualization of c-MYC binding to known target genes. (D) Visualization of c-MYC binding to intergenic regions. (E) Visualization of c-MYC binding to the *Hsf1* locus, revealed by ChIP-seq.

### Figure S3

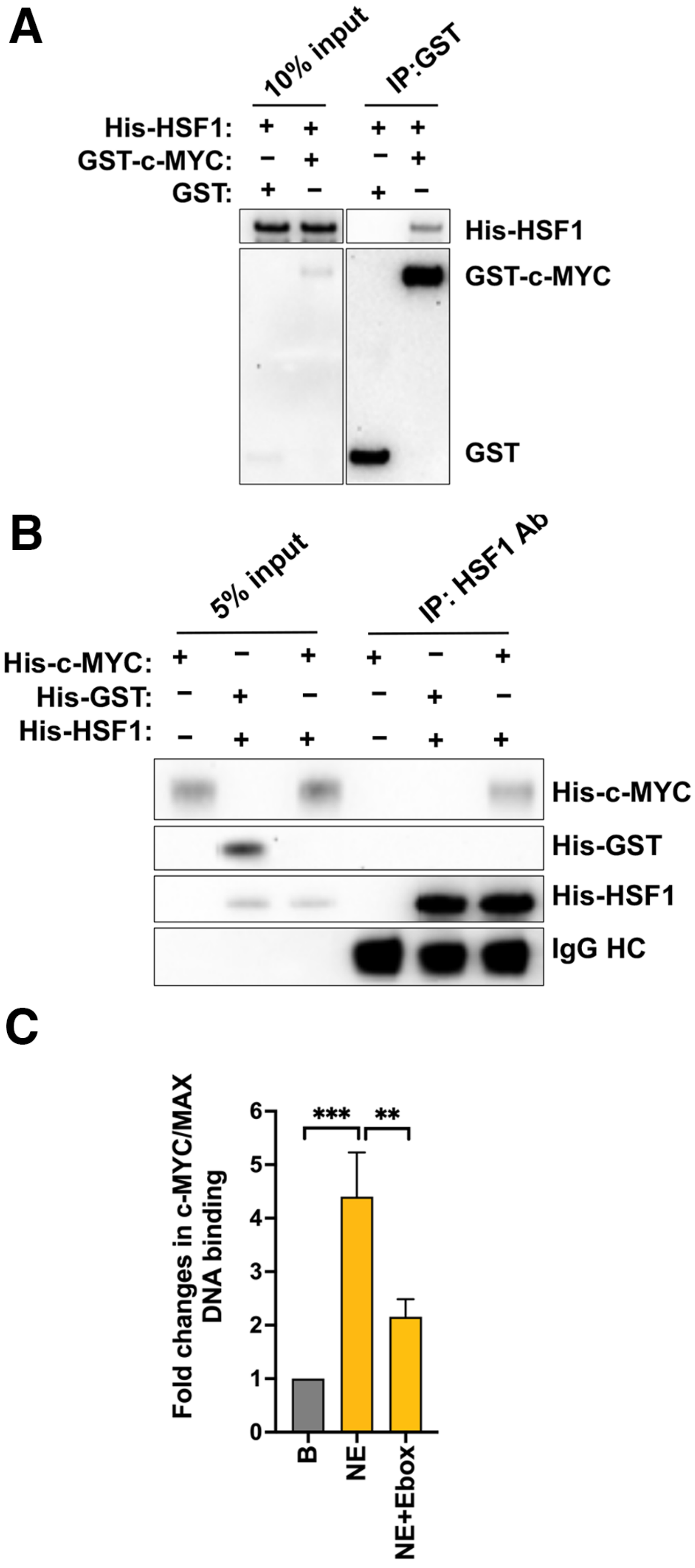
A
10% input
IP:GST
His-HSF1:
GST-c-MYC:
GST:
His-HSF1
GST-c-MYC
GST
B
5% input
IP: HSF1 Ab
His-c-MYC:
His-GST:
His-HSF1:
His-c-MYC
His-GST
His-HSF1
IgG HC
C
Fold changes in c-MYC/MAX DNA binding
***
**
0
1
2
3
4
5
6
B
NE
NE+Ebox

**Figure S3: HSF1 physically interacts with c-MYC.**

(A) *In vitro* pull-down assays were performed with recombinant GST-c-MYC and His-HSF1 proteins using glutathione magnetic beads. (B) *In vitro* pull-down assays were performed with recombinant His-HSF1 and His-c-MYC proteins using a rabbit anti-HSF1 (H-311) antibody. (C) Validation of the *in vitro* c-MYC DNA binding assay. Each well was incubated with 50 µg nuclear extracts with and without the competition of free wild-type E-box oligos (mean ± SD, n =3 independent experiments, One-way ANOVA). B: blank (lysis buffer only). NE: nuclear extracts.

### Figure S4

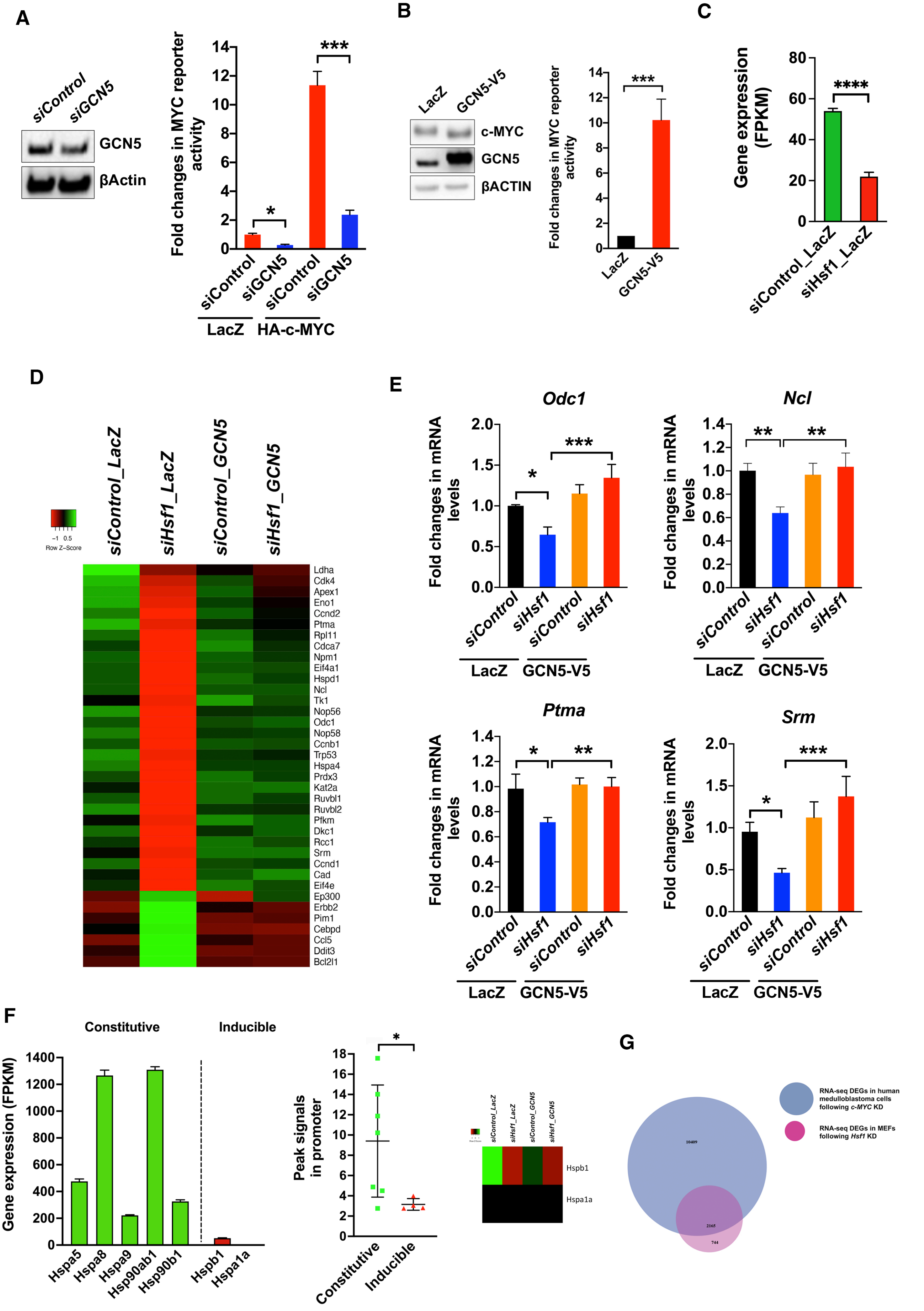

A
siControl
siGCN5
GCN5
βActin
Fold changes in MYC reporter activity
14
12
10
8
6
4
2
0
*
***
siControl
siGCN5
siControl
siGCN5
LacZ
HA-c-MYC
B
LacZ
GCN5-V5
c-MYC
GCN5
βACTIN
Fold changes in MYC reporter activity
***
C
Gene expression (FPKM)
80
60
40
20
0
****
siControl_LacZ
siHsf1_LacZ
D
siControl_LacZ
siHsf1_LacZ
siControl_GCN5
siHsf1_GCN5
Row Z-Score
Ldha
Cdk4
Apex1
Eno1
Ccnd2
Ptma
Rpl11
Cdca7
Npm1
Eif4a1
Hspd1
Ncl
Tk1
Nop56
Odc1
Nop58
Ccnb1
Trp53
Hspa4
Prdx3
Kat2a
Ruvbl1
Ruvbl2
Pfkm
Dkc1
Rcc1
Srm
Ccnd1
Cad
Eif4e
Ep300
Erbb2
Pim1
Cebpd
Ccl5
Ddit3
Bcl2l1
E
Odc1
Ncl
Ptma
Srm
Fold changes in mRNA levels
siControl
siHsf1
siControl
siHsf1
LacZ
GCN5-V5
F
Constitutive
Inducible
Gene expression (FPKM)
1400
1200
1000
800
600
400
200
0
Hspa5
Hspa8
Hspa9
Hsp90ab1
Hsp90b1
Hspb1
Hspa1a
Peak signals in promoter
Constitutive
Inducible
siControl_LacZ
siHsf1_LacZ
siControl_GCN5
siHsf1_GCN5
Hspb1
Hspa1a
G
RNA-seq DEGs in human medulloblastoma cells following c-MYC KD
RNA-seq DEGs in MEFs following Hsf1 KD
10409
2165
744

**Figure S4: HSF1 activates c-MYC via GCN5.**

(A) Left panel: GCN5 protein levels were detected by immunoblotting in HEK293T cells transfected with siRNAs. Right panel: c-MYC transcriptional activities were measured by the dual reporter system in HEK293T cells transfected with indicated plasmids and siRNAs (mean ± SD, n = 3 independent experiments, One-way ANOVA). (B) Left panel: c-MYC protein levels were detected by immunoblotting in immortalized MEFs stably expressing LacZ or V5-GCN5. Right panel: endogenous c-MYC transcriptional activities were measured by the dual reporter system in these MEFs (mean ± SD, n = 3 independent experiments, Two-tailed Student's t test). (C) Quantitation of *Hsf1* mRNA levels by RNA-seq (mean ± SD, n = 3 biological replicates, Two-tailed Student's t test). (D) Heatmap visualization of the expression changes of known c-MYC targets in immortalized MEFs, revealed by RNA-seq (each data point represents the average of three biological replicates). (E) The expression of selective c-MYC target genes were quantitated by qRT-PCR (mean ± SD, n = 3 independent experiments, One-way ANOVA). (F) Left panel: RNA-seq quantitation of the expression of *Hsp* genes in the *siControl*_LacZ MEFs (mean ± SD, n = 3 biological replicates). Middle panel: peak signals at the promoters of *Hsp* genes that are constitutively or inducibly expressed (mean ± SD, n =7 or 4 binding sites, Two-tailed Student's t test). Right panel: heatmap visualization of the gene expression changes of *Hspb1* and *Hspa1a* (each data point represents the average of three biological replicates). (G) Venn diagram showing the overlaps of DEGs between human medulloblastoma cells following *c-MYC* KD and immortalized MEFs following *Hsf1* KD.

### Figure S5

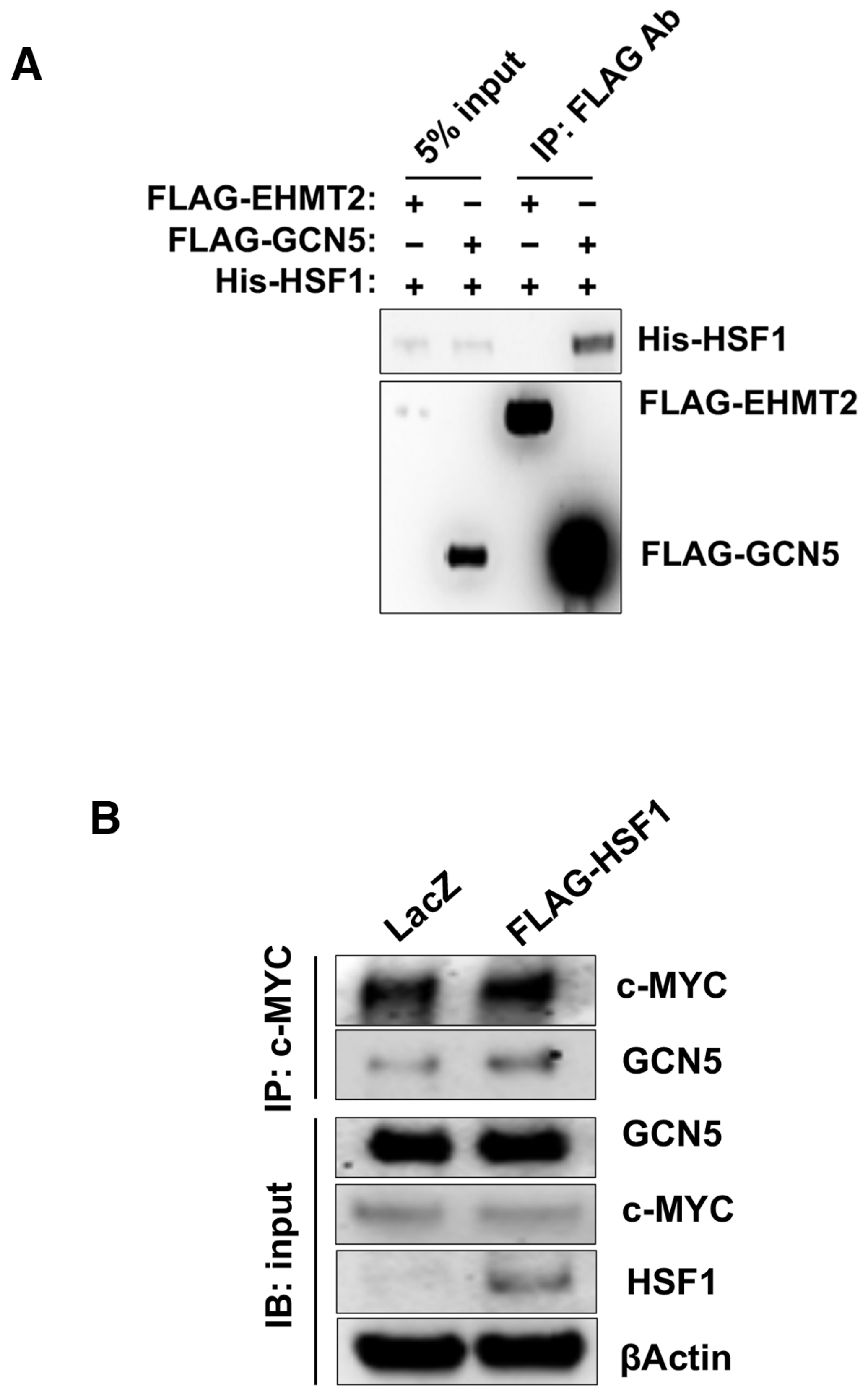
A
5% input
IP: FLAG Ab
FLAG-EHMT2: + – + –
FLAG-GCN5: – + – +
His-HSF1: + + + +
His-HSF1
FLAG-EHMT2
FLAG-GCN5
B
LacZ
FLAG-HSF1
IP: c-MYC
c-MYC
GCN5
IB: input
GCN5
c-MYC
HSF1
βActin

**Figure S5: HSF1 recruits GCN5 to c-MYC.**

(A) *In vitro* pull-down assays were performed with recombinant FLAG-GCN5 and His-HSF1 proteins. FLAG-EHMT2 served as the negative control. (B) Co-IP of endogenous c-MYC and GCN5 in A2058 human melanoma cells stably expressing FLAG-HSF1.

Figure S6

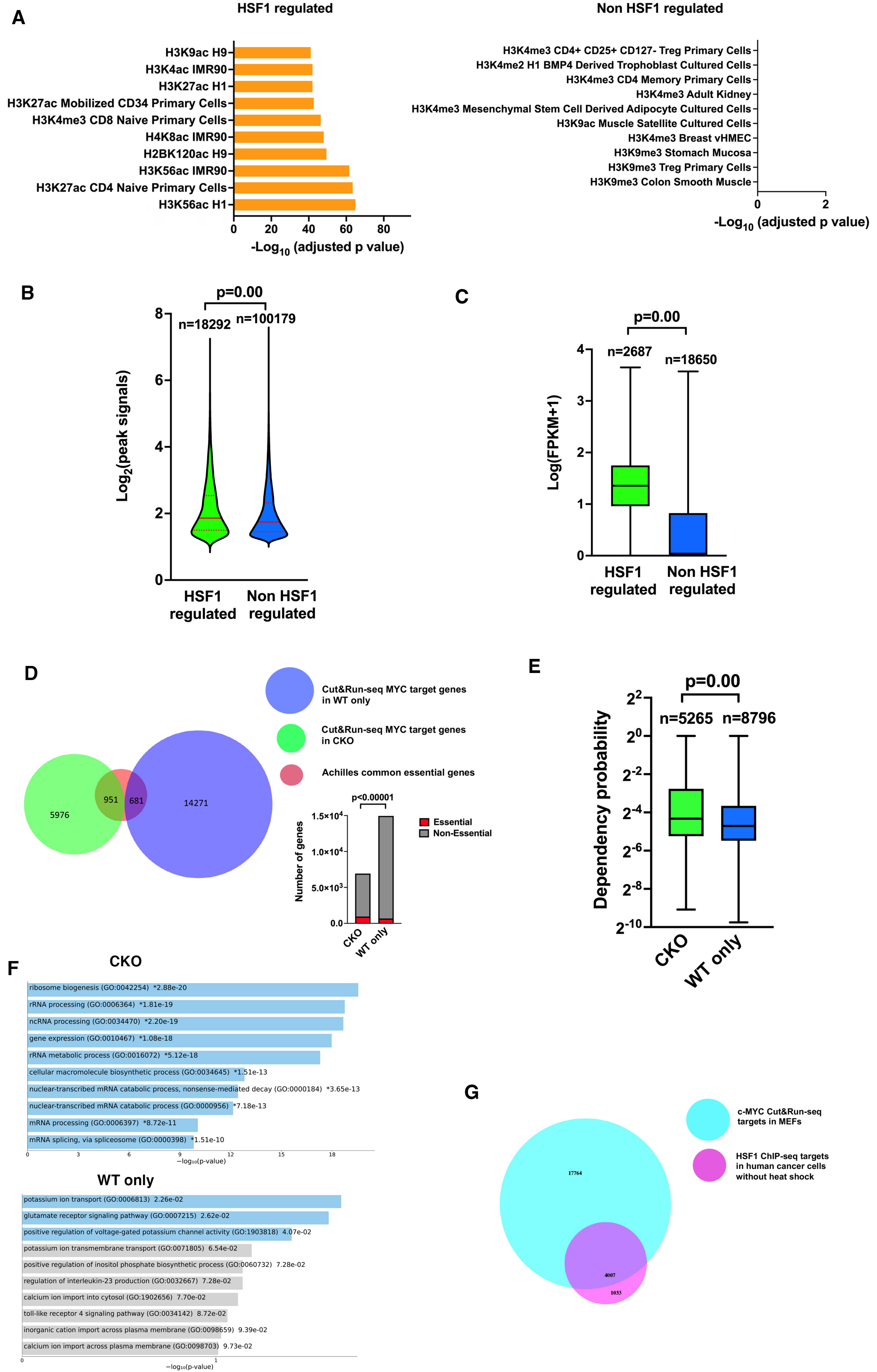

A
HSF1 regulated
H3K9ac H9
H3K4ac IMR90
H3K27ac H1
H3K27ac Mobilized CD34 Primary Cells
H3K4me3 CD8 Naive Primary Cells
H4K8ac IMR90
H2BK120ac H9
H3K56ac IMR90
H3K27ac CD4 Naive Primary Cells
H3K56ac H1
0 20 40 60 80
-Log10 (adjusted p value)
Non HSF1 regulated
H3K4me3 CD4+ CD25+ CD127- Treg Primary Cells
H3K4me2 H1 BMP4 Derived Trophoblast Cultured Cells
H3K4me3 CD4 Memory Primary Cells
H3K4me3 Adult Kidney
H3K4me3 Mesenchymal Stem Cell Derived Adipocyte Cultured Cells
H3K9ac Muscle Satellite Cultured Cells
H3K4me3 Breast vHMEC
H3K9me3 Stomach Mucosa
H3K9me3 Treg Primary Cells
H3K9me3 Colon Smooth Muscle
0 2
-Log10 (adjusted p value)
B
p=0.00
n=18292
n=100179
Log2(peak signals)
0 2 4 6 8
HSF1 regulated
Non HSF1 regulated
C
p=0.00
n=2687
n=18650
Log(FPKM+1)
0 1 2 3 4
HSF1 regulated
Non HSF1 regulated
D
Cut&Run-seq MYC target genes in WT only
Cut&Run-seq MYC target genes in CKO
Achilles common essential genes
5976
951
681
14271
p<0.00001
Number of genes
0.0
5.0×10^3
1.0×10^4
1.5×10^4
CKO
WT only
Essential
Non-Essential
E
p=0.00
n=5265
n=8796
Dependency probability
2^2 2^0 2^-2 2^-4 2^-6 2^-8 2^-10
CKO
WT only
F
CKO
ribosome biogenesis (GO:0042254) *2.88e-20
rRNA processing (GO:0006364) *1.81e-19
ncRNA processing (GO:0034470) *2.20e-19
gene expression (GO:0010467) *1.08e-18
rRNA metabolic process (GO:0016072) *5.12e-18
cellular macromolecule biosynthetic process (GO:0034645) *1.51e-13
nuclear-transcribed mRNA catabolic process, nonsense-mediated decay (GO:0000184) *3.65e-13
nuclear-transcribed mRNA catabolic process (GO:0000956) *7.18e-13
mRNA processing (GO:0006397) *8.72e-11
mRNA splicing, via spliceosome (GO:0000398) *1.51e-10
0 3 6 9 12 15 18
−log10(p-value)
WT only
potassium ion transport (GO:0006813) 2.26e-02
glutamate receptor signaling pathway (GO:0007215) 2.62e-02
positive regulation of voltage-gated potassium channel activity (GO:1903818) 4.07e-02
potassium ion transmembrane transport (GO:0071805) 6.54e-02
positive regulation of inositol phosphate biosynthetic process (GO:0060732) 7.28e-02
regulation of interleukin-23 production (GO:0032667) 7.28e-02
calcium ion import into cytosol (GO:1902656) 7.70e-02
toll-like receptor 4 signaling pathway (GO:0034142) 8.72e-02
inorganic cation import across plasma membrane (GO:0098659) 9.39e-02
calcium ion import across plasma membrane (GO:0098703) 9.73e-02
0 1
−log10(p-value)
G
c-MYC Cut&Run-seq targets in MEFs
HSF1 ChIP-seq targets in human cancer cells without heat shock
17764
4007
1033

**Figure S6: HSF1 potentiates the c-MYC mediated transcription.**

(A) Enrichment analyses of epigenetic marks associated with the HSF1-regulated and non-HSF1-regulated c-MYC target genes. (B) Violin plots of CUT&RUN-seq peak signals at the HSF1-regulated and non-HSF1-regulated c-MYC target genes (solid lines represent medians and dotted lines represent IQR, n=18,292 and 100,179 peaks, Mann-Whitney U test). (C) Box plots of RNA-seq gene expression levels of the HSF1-regulated and non-HSF1-regulated c-MYC target genes. The box bounds the IQR divided by the median and the whiskers extend to the minimum and maximum values (n=2,687 and 18,650 genes, Mann-Whitney U test). (D) Venn diagram showing the overlaps among the Achilles common essential genes, c-MYC target genes in $Hsf1^{CKO}$ MEFs, and c-MYC target genes in $Hsf1^{WT}$ MEFs only. Bar graphs summarize the numbers of essential and non-essential genes in each group (Chi-square test). (E) Box plots of the dependency probability of c-MYC target genes (median, minimum and maximum, n=5,265 and 8,796 genes, Mann-Whitney U test). CKO: target genes in $Hsf1^{CKO}$ MEFs; WT only: target genes in $Hsf1^{WT}$ MEFs only. The Achilles gene dependency data are downloaded from DepMap, Broad (2021): DepMap 21Q4 Public. figshare. Dataset. https://doi.org/10.6084/m9.figshare.16924132.v1. (F) Gene ontology enrichment analysis of the c-MYC target genes in $Hsf1^{CKO}$ MEFs and in $Hsf1^{WT}$ MEFs only. (G) Venn diagram showing the overlap between the c-MYC CUT&RUN-seq target genes in immortalized MEFs and the HSF1 ChIP-seq target genes in human cancer cells without heat shock.
